# Epigenetic regulation of unique genes and repetitive elements by the KRAB zinc finger protein ZFP57

**DOI:** 10.1101/611400

**Authors:** Hui Shi, Ruslan Strogantsev, Nozomi Takahashi, Anastasiya Kazachenka, Matthew C. Lorincz, Myriam Hemberger, Anne C. Ferguson-Smith

## Abstract

**Background:** KRAB-zinc finger proteins (KZFPs) represent one of the largest families of DNA binding proteins in vertebrate genomes and appear to have evolved to silence transposable elements (TEs) including endogenous retroviruses through sequence-specific targeting of repressive chromatin states. ZFP57 is required to maintain the post-fertilization DNA methylation memory of parental-origin at genomic imprints along with ZFP445 which is specific for imprints. However, ZFP57 has multiple methylated genomic targets. Here we conduct RNA-seq and ChIP-seq analyses in normal and ZFP57 mutant mouse ES cells to understand the relative importance of ZFP57 at unique and repetitive regions of the genome.

**Results:** Over 80% of ZFP57 targets are TEs, however, ZFP57 is not essential for their repression. The remaining targets lie within unique imprinted and non-imprinted sequences. Though loss of ZFP57 influences imprinted genes as expected, the majority of unique gene targets lose H3K9me3 with little effect on DNA methylation and very few exhibiting alterations in expression. Comparison with DNA methyltransferase-deleted ES cells (TKO) identifies remarkably similar losses of H3K9me3 and changes in expression, defining regions where H3K9me3 is secondary to DNA methylation. We show that ZFP57 is the principal methylation-sensitive KZFP recruiting KAP1 and H3K9me3 in ES cells. Finally, like imprints, other unique targets of ZFP57 are enriched for germline-derived DNA methylation including oocyte-specific methylation that is resistant to post-fertilisation epigenetic reprogramming.

**Conclusion:** Our analyses suggest the evolution of a rare DNA methylation-sensitive KZFP that is not essential for repeat silencing, but whose primary function is to maintain DNA methylation and repressive histone marks at germline derived imprinting control regions.

## Main Text

Kruppel-associated (KRAB)-zinc finger proteins (KZFPs) represent one of the largest families of DNA binding proteins. They are represented in most but not all vertebrate species [1–4], with a recent study mapping the target sites of over 200 KZFPs in human HEK293T cells predominantly to transposable elements including endogenous retroviruses (ERVs) [4]. This is consistent with a small number of more focused studies investigating individual KZFPs in the regulation of transposable elements. For example, KZFPs such as ZFP809 [5], ZFP932 and its paralog Gm15446 [6], and ZNF91/93 [7] have each been shown to repress different retrotransposon families in mouse embryonic stem cells (ES cells). Collectively, these and other studies [8,9] suggest that the main function of KZFPs is to regulate transposable elements. The binding of KZFPs to retrotransposons is associated with their transcriptional silencing and is mediated by recruitment of the KAP1 (also known as TRIM28 or TIF1β) co-repressor complex, which induces the local acquisition of H3K9me3. Members of the co-repressor complex include the histone methyltransferase SETDB1 (also known as ESET), heterochromatin protein 1 (HP1), the histone demethylase LSD1, and NuRD histone deacetylase complex [10,11]. Loss of KAP1 binding leads to the specific loss of H3K9me3 [12].

Individual KZFPs have also been noted to bind to unique regions of the mammalian genome [4,13–15]. In particular, a role for ZFP57 in the targeted maintenance of genomic imprints has provided novel insights into functions of this family outside the management of the repeat genome. Genomic imprinting is a process that causes genes to be expressed according to their parental origin. Imprinted genes are regulated by parental origin-specific DNA methylation at imprinting control regions (ICRs), that is acquired at different locations in the male and female germlines (germline differentially methylated regions, gDMRs) and maintained after fertilization during preimplantation epigenetic reprogramming [16]. *In vivo*, ZFP57, with ZFP445, is required to maintain the methylation memory of parental-origin during this critical dynamic epigenetic period. ZFP57 binds to all ICRs in ES cells [15,17–20] with the exception of *Slc38a4*, which appears to be regulated in a different manner [21]. In humans, mutation of ZFP57 has been found in patients with multi-locus imprinting disturbances including transient neonatal diabetes mellitus type 1 (TNDM1) [22,23]. Furthermore, in mice, ZFP57 exerts a maternal-zygotic effect, whereby deletion of both the maternal gene in oocytes and the zygotic copies in early embryos causes severe loss of methylation imprints at multiple imprinted loci, resulting in embryonic lethality. Homozygous deletion of only the zygotic ZFP57 presents a partially penetrant lethal phenotype [15]. Binding of ZFP57 to other unique regions has also been described, including strain-specific interactions conferred by genotype specific polymorphisms in ZFP57 recognition sites [18]. Though correlated with strain-specific transcriptional behavior in some instances [18,24], the functional importance of such interactions at non-imprinted loci *in vivo* are not known.

In order to determine the relative importance of ZFP57 binding at different genomic locations, we assessed whether ZFP57 function in ES cells extends beyond the regulation of imprinted regions including at retrotransposons. We found extensive ZFP57 binding at unique germline methylated sites outside imprinted domains, whilst the vast majority of binding was targeted to transposable elements (TEs). Deletion of ZFP57 resulted in loss of KAP1 binding and H3K9me3 at imprinted locations as well as at ∼100 other unique ZFP57 targets. In contrast to imprinted domains, this was only modestly correlated with changes in expression of nearby genes suggesting a functionally distinct role for ZFP57 at these regions. With few notable exceptions, TEs maintained H3K9me3 supporting the idea that they may be transcriptionally silenced by multiple redundant mechanisms involving other KZFPs in ES cells [4,6,25,26]. This pattern of retained H3K9me3 was also observed in ES cells derived from embryos with maternal-zygotic ZFP57 deletion. Interestingly, the pattern of ZFP57 binding in ES cells essentially mimicked sites where H3K9me3 was lost in DNA methyltransferase (*Dnmt1/Dnmt3a/Dnmt3b*) triple knock out (DNMT TKO) ES cells [26], indicating a primary role for DNA methylation in the establishment of H3K9me3 via ZFP57. Thus, we propose that the primary role of ZFP57 is in maintenance of genomic imprints via its DNA methylation-sensitive binding, and that its interaction with other targets occurs by virtue of their existing DNA methylation status, perhaps providing reinforcement of an already silenced state.

## Results

### Analysis of blastocyst derived ZFP57 null ES cells

To better understand the role(s) played by ZFP57 during preimplantation development we have derived ZFP57 null ES cells directly from C57BL/6J (BL6) blastocyst embryos in ground state pluripotency conditions [27]. ZFP57 homozygous mutant ES cells (ZFP57 KO) were derived from embryos which targeted coding exon 4 and 5 comprising both the KRAB domain and zinc fingers [19]. Genotyping analysis of ES cells showed the expected Mendelian ratios, enabling selection of both wild-type and mutant male lines for further analysis. In addition, we derived two maternal-zygotic ZFP57 deleted (ZFP57 MZ-KO) ES cell lines on mixed C57BL/6J–129/SvS1 (BL6/129) background. These could not be generated on a pure C57BL/6J background since homozygous BL6 mutant females do not survive to term. Absence of ZFP57 protein in mutants was confirmed by western blot analysis on whole cell extracts (Fig. S1A). Quantitative bisulfite pyrosequencing confirmed loss of methylation imprints in all the mutants consistent with previous reports [17,24], with the exception of KvDMR1, where no loss of DNA methylation was observed in these cells (Fig. S1B). These results show for the first time, that loss of imprinting is equivalent in both zygotic and maternal-zygotic ZFP57 mutant ES cells.

### Mapping of genome-wide ZFP57 binding sites in ES cells reveals binding events at transposable elements in addition to unique non-imprinted genes

We mapped ZFP57 genomic targets by ChIP-seq in WT and ZFP57 KO ES cell lines using an antibody that recognizes endogenous ZFP57. In total, we identified 1423 high-confidence ZFP57 peaks, where each peak was independently called in at least two of three WT biological replicates. As expected, we observed strong ZFP57 binding at 20 of the 22 known ICR elements, including binding at previously identified tissue specific imprinted genes *Cdh15* [28], with weaker peaks at the two remaining imprints lying below the peak threshold (*Gpr1* and *Mcts2*). As in our previous report, no binding was detected at the *Slc38a4* DMR [18], in agreement with study demonstrating a role for H3K27me3 in regulating its germline imprinting [21].

Using this stringent cut-off, the remainder of the ZFP57 peaks were further characterized according to their relative location at unique regions and at different types of repetitive elements in the mouse genome. We found 13.5% (n=189) targets mapped to non-repetitive regions, a further small subset of peaks 1.3% (n=19) at non-transposon derived repeats (e.g. Simple, low complexity and satellite repeats), whilst the vast majority of the peaks 85.2% (n=1194) targeted transposable elements (Fig. 1A). In total, ZFP57 binds 1061 LTR containing retrotransposons of which the majority (n=795) were IAPs, with IAPLTR2a2_Mm representing the largest class (Fig. 1B). In total, ∼7-8% of all IAPs present in the mouse genome are bound by ZFP57 in ES cells. This binding to repeats was highly specific since only uniquely aligned reads were used for peak calling (based on paired-end read mapping) (Fig. S2). TE enrichment in the ZFP57-bound subset of repeats was highly significant in comparison to the rest of genome (Fig. 1C, *P* < 0.0001).

**Fig. 1.**
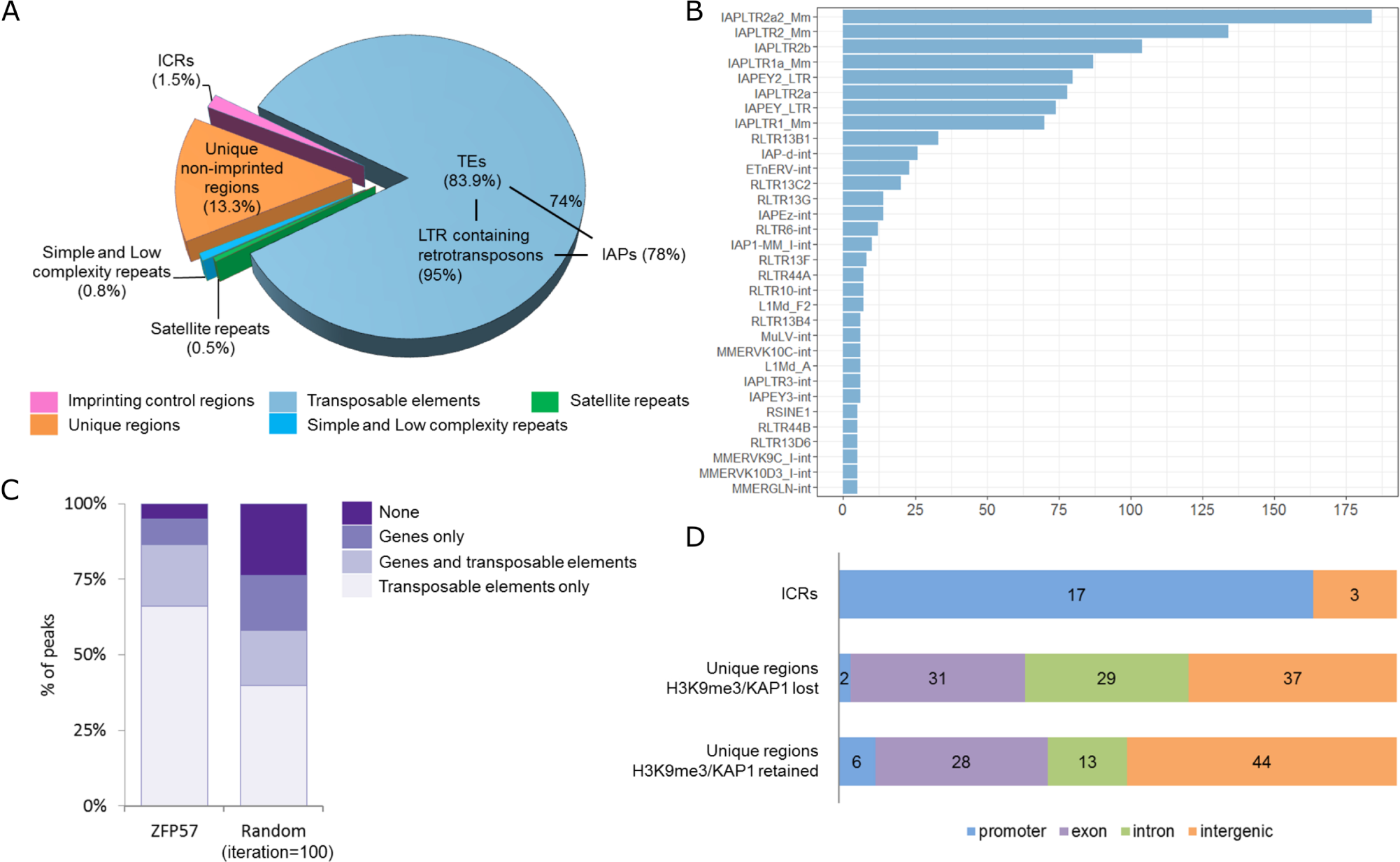
Genome-wide mapping of ZFP57 binding sites in ES cells. (A) Distribution of ZFP57 binding sites with respect to unique regions and repetitive elements. (B) Top 32 TE classes bound by ZFP57, ranked by number of ZFP57 peaks. IAPLTR2a2_Mm represents the most commonly targeted TE by ZFP57. (C) Significant enrichment of ZFP57 peaks at TEs relative to random peak permutations, number of iterations =100, binomial *P* < 0.0001. (D) Non-repeat associated peaks bind distinct genomic regions within ICRs and non-imprinted regions.

Analysis of non-repeat associated ZFP57 peaks revealed a distinct distribution of imprinted versus non-imprint bound sites with respect to gene features. Whilst maternally methylated ICRs coincide with gene promoters, the majority of non-imprint associated peaks were equally distributed within gene exons, introns and intergenic regions (Fig. 1D). Indeed, very few promoter-bound sites lay outside imprinted regions, but notably included the promoter of the *Dux* gene – a transcription factor involved in zygotic genome activation at the 2 cell stage [29,30]. Our analysis of data from Imbeault et al, revealed that the human DUX4 promoter is targeted by multiple KZFPs including ZFP57 [4], consistent with our observation that H3K9me3 is only partially lost at this locus in ZFP57 mutants.

### Distinct H3K9me3 and KAP1 binding profile at ZFP57 bound unique regions and transposable elements

KZFPs are associated with the KAP1 mediated recruitment of SETDB1 and in turn H3K9me3 deposition. We therefore examined the presence or absence of KAP1 and H3K9me3 in ZFP57 KO cells by ChIP-seq. Once again, using ICRs as internal controls, we found the expected loss of KAP1 binding at almost all ICRs (n=20), of which 17 also exhibited complete loss of H3K9me3 (Fig. 2 and Fig. S3). Interestingly, H3K9me3 was also lost at *KvDMR1* ICR – the only DMR retaining DNA methylation in the KO ES cells (Fig. S1a and Fig. S3). Three ICRs: *Grb10, Fkbp6* and *Peg10* exhibited varying degrees of loss of H3K9me3 and KAP1 binding (Fig. S4).

**Fig. 2.**
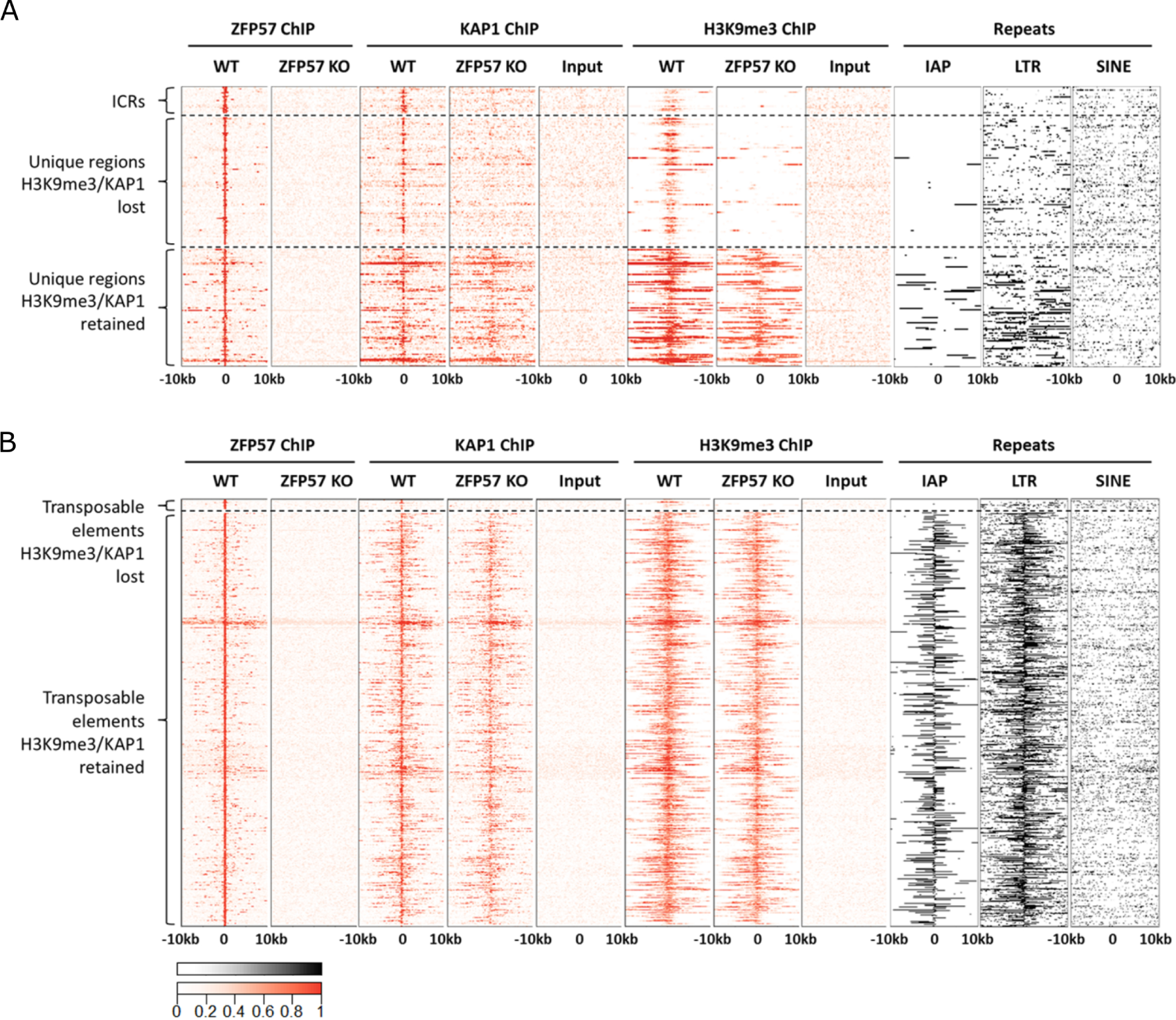
Distinct H3K9me3 and KAP1 binding profile at ZFP57 bound unique regions and transposable elements. (A) Heat maps of ChIP-seq enrichment signals (red) within ICRs/unique regions and (B) transposable elements associated binding sites. Enrichment of IAP, LTR and SINE elements are shown as black-and-white heat maps. Within unique regions, significant enrichment for LTRs nearby H3K9me3 retaining group compared to H3K9me3 losing group was observed by measuring distance to the nearest LTR from each peak (unpaired two-tailed t-test, *P* < 0.001).

We next analyzed the genome-wide distribution of KAP1 and H3K9me3 at different categories of ZFP57 peaks in WT and KO cells. Approximately one half (n=98, 52%) of the unique non-ICR peaks exhibited complete loss of KAP1 and H3K9me3 in ZFP57 mutant ES cells, whilst the rest (n=91, 48%) retained KAP1 binding and H3K9me3 (Fig. 2A). Upon further inspection of the H3K9me3 retaining group it was revealed that although mapping to unique sequences, they are found in more repeat-dense genomic regions with specific enrichment of LTR-containing retrotransposons (*P* < 0.001, unpaired two-tailed t-test) encompassing IAPs (Fig. 2A, right panel). We speculate that KAP1 binding (presumably recruited by other KZFPs) at those nearby retrotransposons maintains a generally repressed state of the entire region irrespective of ZFP57 binding.

In contrast to the unique ZFP57 target sites, virtually all ZFP57 peaks at transposable elements (98%; n=1168) retained H3K9me3 levels (Fig. 2B). Only 26 (2.2%) TEs showed loss of KAP1 and H3K9me3 and these were mainly SINEs, non-IAP retroelements or solo IAP-derived LTRs. We thus conclude that despite widespread binding of ZFP57 to transposable elements, it is not required for their repression.

Finally, we mapped H3K9me3 profiles in ES cells derived from maternal-zygotic deletion of ZFP57 (Fig. S5). Despite the more penetrant phenotype *in vivo* of the maternal zygotic deletion [15], we observed a nearly identical pattern of H3K9me3 loss/retention over the unique and repeat associated subsets of peaks in ZFP57 MZ-KO ES cells.

### Transcriptional profiling identifies few specific non-imprinted ZFP57 target genes

To determine the functional consequences of ZFP57 deletion on the transcriptome, total RNA sequencing post-ribosomal RNA (rRNA) depletion was performed on these same control and mutant cell lines.

Genome-wide differential expression analysis revealed statistically significant altered expression of 1080 genes in ZFP57 KO cells in comparison to WT, which included many known imprinted genes in agreement with previously demonstrated roles of ZFP57 in imprint regulation (Fig. 3A). Quantification of all annotated gene transcripts followed by hierarchical clustering analysis indicated clear segregation of the WT and mutant cell lines (Fig. 3B). In order to determine whether non-imprint bound ZFP57 regions might regulate expression of nearby genes, we analyzed expression of all genes in close proximity (< 20 kb) to ZFP57 peaks (n=748, Fig. 3C). We found that only a small subset (n=75) of differentially expressed genes harbored ZFP57 binding sites or have ZFP57 peaks located nearby (< 20kb). Thus, the majority of differentially expressed genes may not be direct targets of ZFP57 and instead may represent secondary effects of a perturbed imprinted gene network.

**Fig. 3.**
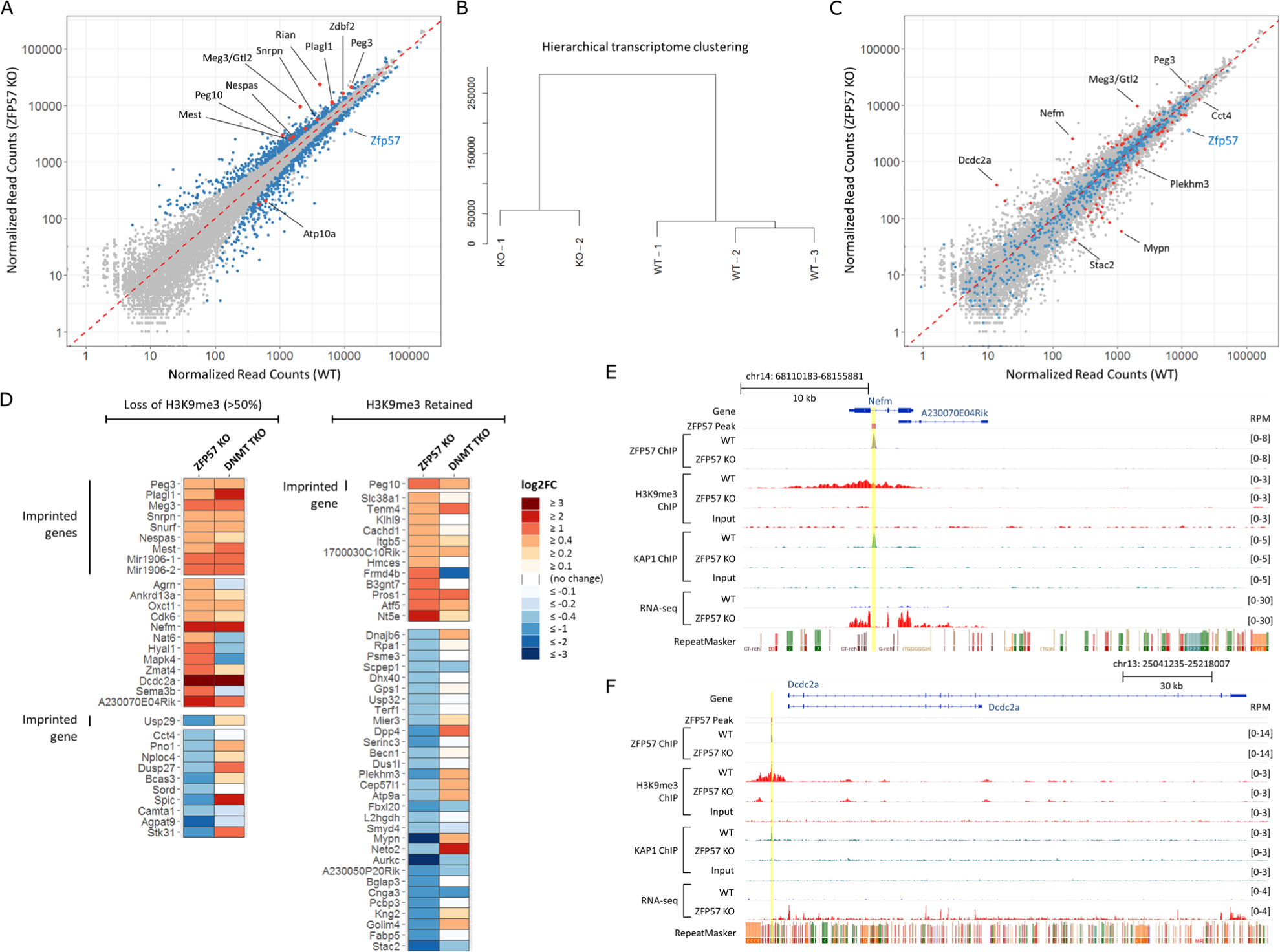
ZFP57 KO ES cells show perturbed expression of imprinted and non-imprinted genes. (A) Scatterplot of gene expression in WT and ZFP57 KO ES cells with differentially expressed genes identified by DESeq2 shown in blue, of which the known imprinted genes further highlighted in red. (B) Hierarchical transcriptome clustering analysis showing clear segregation of WT versus KO cells. (C) Scatterplot as in (A), but highlighted all the genes < 20kb of ZFP57 peak, blue = no significant change in expression of associated genes (n=673), red = differentially expressed (n=75). (D) Gene expression differences within the 75 genes, with ZFP57 peaks losing H3K9me3 shown on the left, and those retaining H3K9me3 shown on the right. Expression of ZFP57 KO and DNMT TKO versus their corresponding WT control is shown side by side for comparison. (E) Genome browser screenshots for the two highest upregulated non-imprinted genes Nefm and (F) Dcdc2a, showing relative location of ZFP57 peak, H3K9me3/KAP1 enrichment and RNA-seq tracks for each gene.

We ascertained whether the 75 ZFP57-bound differentially expressed genes were misregulated as a direct consequence of loss of H3K9me3 and/or DNA methylation in ZFP57 KO ES cells by comparing this with their expression levels in DNMT TKO ES cells lacking both maintenance and de novo DNA methyl transferases [26]. These cells are devoid of any detectable 5-methylcytosine (5mC) in the genome, and as ZFP57 binding is DNA methylation dependent, they are unable to target ZFP57 binding. We found that genes upregulated in ZFP57 KO ES cells had a strong tendency to be also upregulated in DNMT TKO cells suggesting these are direct ZFP57 targets (Fig. 3D, top). This correlation was strongest amongst imprinted genes. In contrast, no such correlation could be seen in genes that are downregulated in ZFP57 null ES cells (Fig. 3D, bottom) indicating that those are likely to be secondary effects. Amongst the strongest upregulated non-imprinted genes in both mutants were the *Nefm* and *Dcdc2a* genes (Fig. 3E and Fig. 3F). In both cases, loss of ZFP57 led to loss of KAP1 binding, and abolition of a repressive H3K9me3 domain around the gene indicating a primary role for DNA methylation in the recruitment of repressive epigenetic state at these regions via ZFP57.

In conclusion, we find a large network of gene deregulation occurring in ZFP57 KO ES cells, however, apart from imprints, only a small proportion of non-imprinted genes are likely to be directly regulated by ZFP57.

### ZFP57 at transposable elements

Given the widely accepted role of KZFPs in silencing retroelements, we further explored whether ZFP57 binding to transposable elements is functional or coincidental. The vast majority of ZFP57-bound transposable elements retained H3K9me3 in mutants (Fig. 2B) and remained transcriptionally repressed (Fig. S6). Interestingly, despite the overall retention of H3K9me3 enrichment, we observed a reduction of KAP1 signal at ZFP57 bound TEs in the mutants (Fig. 4A), indicating some contribution to the recruitment of KAP1. We hypothesized that the retained repression of retrotransposons in ZFP57 KO ES cells might be mediated by binding of other KZFPs recruiting KAP1 at levels sufficient to maintain H3K9me3. A recent screen for KZFPs that target retrotransposon silencing identified ZFP932 and Gm15446 as partially redundant factors binding to endogenous retroviruses including IAPs [6]. We overlapped these published binding sites with those at ZFP57 and identified a subset (n=85) of targeted transposable elements that are jointly bound by ZFP57 (Fig. 4B). One example of this is shown for the RLTR44E retroelement located ∼2kb downstream from the *Ankrd10* gene and harbors adjacently located ZFP57 and ZFP932/Gm15446 peaks (Fig. 4C). In ZFP57 null ES cells, KAP1 is specifically lost under the ZFP57 peak, whilst being maintained at the neighboring site of two other KZFPs. Consequently, the region of the repeat maintains overall H3K9me3 levels and neither reactivation of RLTR44E nor *Ankrd10* was observed in ZFP57 mutants. Amongst triply bound transposable elements in general, no reactivation was observed in either the ZFP932/Gm15446 double deletion or ZFP57 KO ES cells (with exception of the IAP retroelement in the vicinity of *Bglap3*), indicating potential for redundant functions of KZFPs (Fig. S7).

**Fig. 4.**
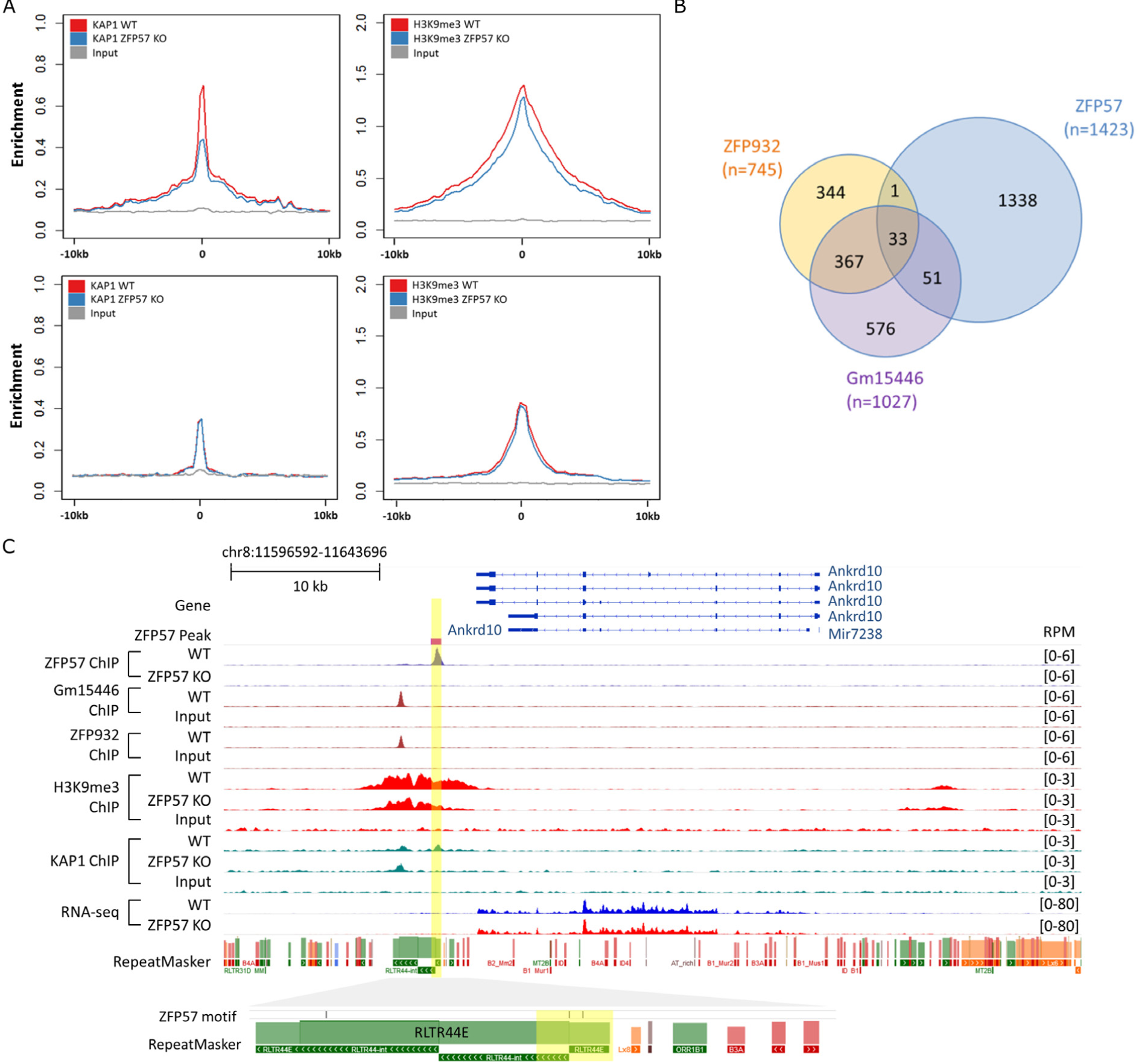
ZFP57 may contribute to transposon silencing. **a**, Reduction of KAP1 ChIP signals in ZFP57 KO under TE-bound ZFP57 peaks, which leads to very modest reduction in H3K9me3 levels (top), no such reduction was observed at other KAP bound TEs in absence of ZFP57 binding (bottom). **b**, Venn diagram showing partially overlapping ZFP57, ZFP932 and Gm15446 peaks. 33 ERVs were found to be triply co-bound. **c**, Genome browser screenshot of a triply co-bound RLTR44E retroelement near Ankrd10 gene. Notably, there’s a partial loss of H3K9me3 under ZFP57 peak in ZFP57 KO but not adjacent ZFP932/Gm15446 peaks, thus maintaining overall silencing of the repeat and unchanged levels of the proximal gene.

Intriguingly, ∼2% of transposable elements (n=26) bound by ZFP57 did exhibit loss of KAP1 and H3K9me3 (Fig. 2B). However, apart from two notable exceptions this was not associated with transcriptional activation of the repeat itself or of the nearby coding gene. In one case, loss of ZFP57 binding at the intergenic MER20 (a DNA transposon) led to an increase in expression of an unannotated transcript comprising both unique regions and a cluster of transposable elements (Fig. S8). In the other example, the IAP derived LTR showed loss of H3K9me3 and KAP1 binding and this was associated with increased expression of the host coding gene *Mapk4* (Fig. S9). Notably this element comprised the LTR only and not the full length IAP, supporting the general rule that IAPs are not affected by ZFP57 ablation.

Together our data suggest that despite widespread binding of ZFP57 at transposable elements, only a very small subset of them is dependent on this to maintain H3K9me3. This might be partially explained by other KZFPs binding alongside ZFP57 to recruit repressive states to these elements. However, the highly redundant and combinatorial nature of KZFPs binding to transposable elements makes it difficult to ascertain the complexities of these relationships.

### Relationship between DNA methylation and H3K9me3 at ZFP57 targets

In order to explore the relationship between DNA methylation and H3K9me3 at different types of ZFP57 targets in more detail, we analyzed DNA methylation levels at these regions using quantitative bisulfite pyrosequencing. As expected, we found that loss of ZFP57 and H3K9me3 was associated with loss of DNA methylation at imprints. Regions retaining H3K9me3 (both transposons and unique non-imprinted peaks) remain DNA methylated (Fig. 5A). Interestingly, non-imprinted regions that lost H3K9me3 retained considerable levels of DNA methylation indicating that DNA methylation at these loci was not dependent on H3K9me3.

**Fig. 5.**
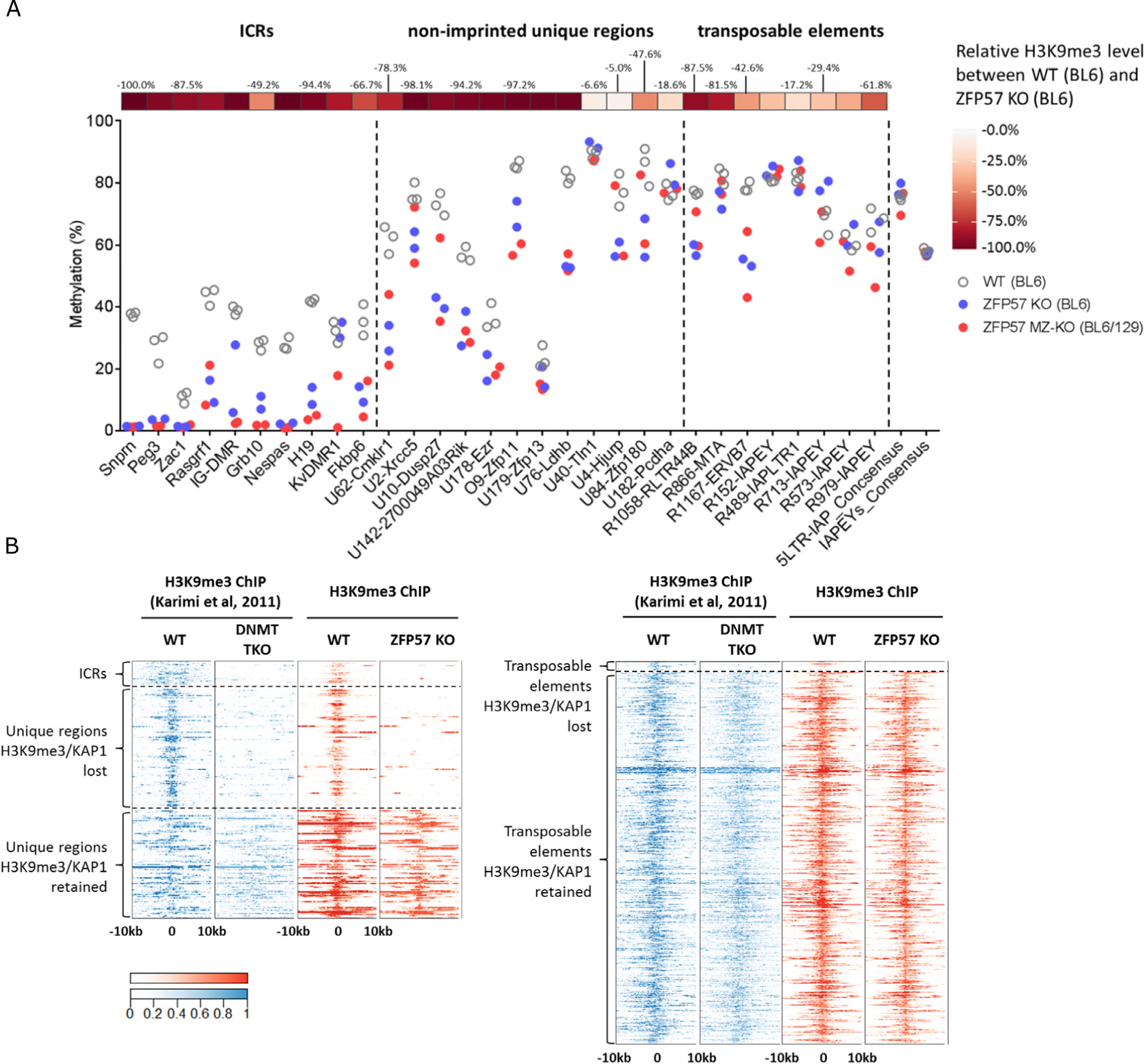
Relationship between DNA methylation and H3K9me3 at ZFP57 binding sites in ES cells. (A) Relationship between loss of DNA methylation and loss of H3K9me3 at select ICRs, unique regions and transposable elements bound by ZFP57. 5’ LTR-IAP and IAPEY consensus regions were used as control. (B) Heat maps of ChIP signals for H3K9me3 in DNMT TKO (blue trace) and ZFP57 KO (red trace) ES cells at ZFP57 bound unique regions (left) and transposable elements (right).

Conversely, in order to test if DNA methylation is necessary to confer H3K9me3 at these regions, we compared the profile of H3K9me3 in ZFP57 KO cells with that of DNA methylation deficient (DNMT TKO) cells [26]. Strikingly, our analysis identified the pattern of H3K9me3 to be virtually identical between these two mutants: the repressive mark was lost at ICRs and the exact same subset of non-imprinted unique regions, whilst maintained over TE-associated ZFP57 peaks (Fig. 5B). Importantly, these data suggest two functions for ZFP57 - one in the maintenance of methylation at imprinted domains, and a second in the methylation-sensitive recruitment of H3K9me3 to some repressed chromatin domains.

Finally, we queried the global extent to which ZFP57 binding sites can explain DNA methylation-dependent H3K9me3 in DNMT TKO cells. In total 15.6% (n=91) of all DNA methylation labile H3K9me3 loci had a strong ZFP57 and KAP1 enrichment signal (Fig. S10). Curiously, the remaining regions (n=492) did not associate with KAP1 indicating that ZFP57 is the main, if not sole, KZFP that recruits repressive histone marks in a DNA methylation-dependent manner in ES cells and that other KAP1-independent processes regulate H3K9me3 at the other sites.

### ZFP57 maintains germline derived DNA methylation at both imprinted and non-imprinted regions during preimplantation development

Imprinting control regions acquire differential DNA methylation in gametes, which is then selectively maintained through preimplantation stages of development. This maintenance has been shown to be ZFP57-dependent *in vivo* for a subset of ICRs [15,19]. We investigated whether non-imprinted unique ZFP57 targets in ES cells are also germline methylated and whether they too are protected from post-fertilization genome-wide erasure of DNA methylation (Fig. 6).

**Fig. 6.**
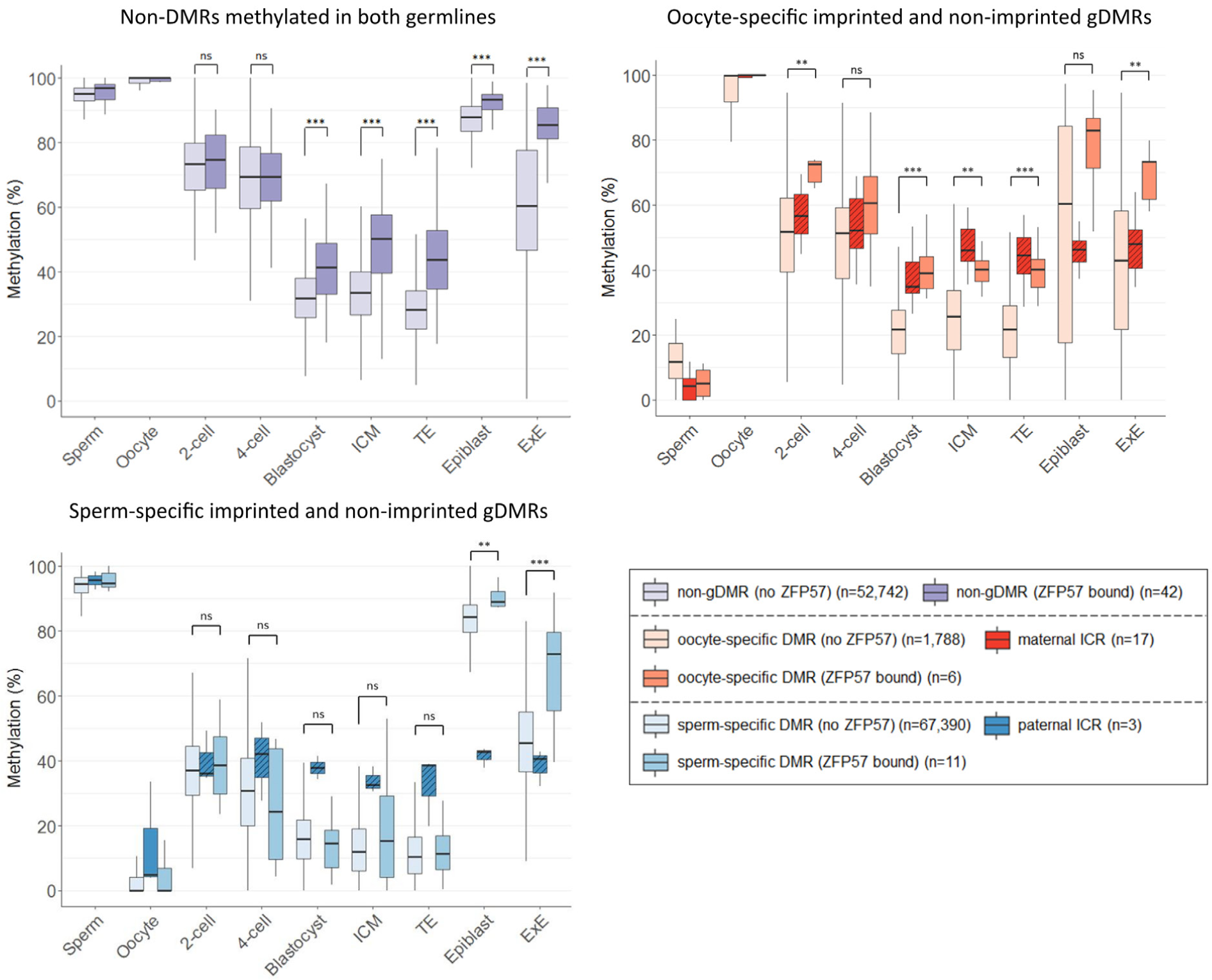
ZFP57 maintains DNA methylation at both imprinted and non-imprinted regions during preimplantation development. DNA methylation level of non-DMRs, oocyte and sperm-specific germline DMRs bound by ZFP57 (darker coloured) versus those not bound by ZFP57 (light coloured) during preimplantation development and in E6.5 epiblast and extraembryonic ectoderm [31–33]. Known ICRs are shown as hatched boxes for comparison. ICM – inner cell mass, TE – trophectoderm, ExE – extraembryonic ectoderm. ns – not significant, * *P* < 0.05, ** *P* < 0.01, *** *P* < 0.001; Mann-Whitney *U* Test.

Analysis of publicly available datasets [31–33] revealed that indeed the majority of unique non-imprinted targets of ZFP57 where methylation data is available are methylated in both germlines (n=42), with a few that are either oocyte (n=6) or sperm (n=11) specific gDMRs. We analysed the DNA methylation dynamics at these regions throughout pre-implantation development and found that ZFP57 binding conferred significant protection against post-fertilization demethylation in all oocyte methylated regions regardless whether they are gDMR or not (Fig. 6). Interestingly, sperm-specific methylated regions with exception of the three known paternal germline imprints are not protected from DNA methylation reprogramming despite being ZFP57 bound in our ChIP-seq data. This hints at potentially distinct mechanisms of maintaining germline-derived DNA methylation on the two parental chromosomes by ZFP57. Oocyte-specific gDMRs bound by ZFP57 resolved their DMR status by acquiring DNA methylation in the post-implantation epiblast, resembling patterns that have been previously reported for placenta-specific imprinted and transient maternal gDMRs [28,33–35]. Though numbers are low, sequence composition analysis revealed that non-imprinted paternal DMRs had a lower CpG density compared to the other groups with the maternal ICRs having the highest CpG percentages (Figure S11).

Taken together, we conclude that ZFP57 can protect against post-fertilization epigenetic reprogramming at sequences methylated in either or both germlines, and such protection may be associated with local CpG density.

## Discussion

In the present study, we demonstrate that in ES cells ZFP57 binds not only to ICRs, but also to non-imprinted unique regions and transposable elements; in particular, to IAPs. We have performed a detailed epigenetic and transcriptomic analysis of ZFP57 mutant ES cells allowing us to ascertain the relative functional roles played by ZFP57 within different genomic locations. We have determined the hierarchical relationship between DNA methylation and H3K9me3 at ZFP57-bound regions, and assessed whether ZFP57 maintains DNA methylation at non-imprinted loci during post-fertilization epigenetic reprogramming.

We have identified a subset of unique non-imprinted genomic regions, which depend on DNA methylation and ZFP57 binding for the recruitment of KAP1 and H3K9me3. Only a small proportion of these were associated with significant changes in gene expression in ZFP57 mutants. A striking difference between ICRs and non-imprinted unique ZFP57-bound regions is that the former are predominantly found at promoters, but the latter bind exonic, intronic and intergenic regions. A subset of ZFP57 peaks mapped to the 3’ exons of several protein coding genes (n=16) including other Zn-finger transcription factors such as *Zfp13, Zfp180* and *Zfp629*. These genes are expressed in WT ES cells and their expression remained unchanged in mutant ES cells despite loss of KAP1 and H3K9me3 over the 3’ end of these genes. Consistent with this, it has been postulated that 3’ end H3K9me3 recruitment may promote genomic stability and prevent recombination between homologous Zn-finger proteins rather than directly regulate their expression [36,37].

DNA methylation and H3K9me3 generally co-localize at repressed genomic regions. However, it is not clear where DNA methylation is the driver for the recruitment of H3K9me3 or where it might occur as a secondary consequence of H3K9me3. Studies to investigate this ‘cause or consequence’ relationship are confounded by the lethality of cells harboring mutations in the H3K9me3 machinery [12,38–40]. Karimi and colleagues have suggested that DNA methylation and H3K9me3 serve to repress distinct sets of genes as well as some classes of retroelements and showed that only a small subset of regions lose their H3K9me3 in DNA methylation deficient (DNMT TKO) ES cells [26]. Comparison of the pattern of H3K9me3 loss in DNMT TKO cells with that in ZFP57 KO cells revealed remarkable similarity. The comparison of H3K9me3 in DNMT TKO and ZFP57KO has therefore identified regions where the presence of H3K9me3 is dependent on DNA methylation and showed that it is ZFP57 bound to methylated DNA that is responsible for the H3K9me3 recruitment. The similarity of the transcriptomes in these two mutant cell types is consistent with this and identifies a key role for ZFP57 in the regulation of transcriptional repression of methylated targets in ES cells. Furthermore, ZFP57 binding sites explain a significant proportion of these DNA-methylation dependent H3K9me3 modified regions, including almost all that are KAP1 dependent. This strongly suggests that ZFP57 is the major KZFP that binds DNA in a methylcytosine-dependent manner in mouse ES cells.

Recently, we showed that another KZFP member, ZFP445 also binds the methylated allele at imprinted gDMRs and *in vivo* contributes to maintenance of DNA methylation at a subset of ICRs along with ZFP57 [20]. However, data presented here suggests that at least in context of mouse embryonic stem cells, ZFP57 is both necessary and sufficient to recruit KAP1 and maintain H3K9me3/DNA methylation at imprints. Furthermore, our analysis of ZFP445 binding sites (data not shown) revealed very little overlap with ZFP57 outside of imprinted regions with TE-binding being a unique feature of ZFP57.

While complete loss of KAP1 and H3K9me3 was observed at ICRs and over half of ZFP57 peaks at unique non-imprinted regions, we found only ∼2% of repeat-associated ZFP57 peaks exhibited complete loss of these marks. Our data support the hypothesis that in mammalian ES cells, retroviral silencing is mediating by multiple factors acting in redundant manner. Indeed, we identified extensive overlaps with two other KZFPs (Zfp932 and Gm15446) previously demonstrated to recruit KAP1/H3K9me3 repressive marks to ERVs including IAPs [6]. In this context it is interesting to consider the DNA methylation-sensitive nature of ZFP57 binding [19,41]. This property makes it an unlikely candidate KZFP for the initiation of silencing at the integration site of ERV retro-transposons since it would require DNA methylation to be established first in order for binding to occur. Rather it may contribute to silencing initiated by other KZFPs such as Zfp809 binding to invariable PBS-regions of IAPs [5,9]. Recent studies have suggested that LTR regions have the potential to function as enhancer and/or alternative promoter elements and can be co-opted by the host genome in regulation of its own gene expression [42–45]. The ability of a KZFP to target repressive chromatin states in a methylation-sensitive manner could provide an additional layer of modulation at such loci. However, we also cannot rule out the possibility that ZFP57 binding at these elements is a functionally irrelevant secondary consequence of DNA methylation occurring at already H3K9me3-repressed LTRs that contain the rather common ZFP57 hexanucleotide binding motif.

We can now speculate about the function and evolution of ZFP57 in the genome. Indeed, we have shown that ZFP57 mutation does not itself influence the silencing of endogenous retroviral sequences but rather conveys DNA methylation protection to both imprinted and non-imprinted unique CpG rich regions methylated in either or both germlines. Such events may result in the altered dosage of nearby genes, which can then be selected for (e.g. in the case of growth regulating genes) and indeed re-enforced by acquiring extra ZFP57 motifs. Indeed, the majority of known imprinted ICRs contain an unusually high number of ZFP57 motifs within otherwise highly CpG-rich regions setting them apart from other CGIs in the genome. The function, if any, of non-imprinted unique ZFP57 bound regions, which apparently have moderate CpG density and one or two motif instances per peak remains to be determined. We therefore propose that ZFP57 evolved from ancestral KZFPs originally acting to manage retrotransposons, but now functions primarily not to silence these repetitive sequences but rather to control genomic imprinting. This is consistent with the finding that ZFP57 also contributes to the regulation of imprinting in humans [23], a species that lacks the CpG rich IAP retrotransposon sequences to which ZFP57 binds in the mouse.

## Data availability

All raw and processed data for ChIP-seq and RNA-seq have been submitted to Gene Expression Omnibus (GEO) database under accession number GSE123942.

## Acknowledgements

This work was funded by grants from the BBSRC (BB/G020930/1) and Wellcome Trust (WT095606) to AFS.

## Authors’ contributions

RS and ACF-S designed the study and conceived the experiments. R.S. and N.T. performed the experiments. H.S. analyzed the data, performed statistical analysis and prepared the figures. A.K. assisted in analysis of repeat data. M.L. provided DNMT TKO data and advised on analysis and data interpretation. H.S., R.S. and A.C.F.-S. interpreted data and wrote the manuscript. M.H. and A.C.F.-S. provided supervision.

## Supplementary Materials

Materials and Methods

Fig. S1-S11

## References

1. Urrutia R KRAB-containing zinc-finger repressor proteins. Genome Biol. 2003;4:231.

2. Huntley S, Baggott DM, Hamilton AT, Tran-Gyamfi M, Yang S, Kim J, et al. A comprehensive catalog of human KRAB-associated zinc finger genes: Insights into the evolutionary history of a large family of transcriptional repressors. Genome Res. 2006;16:669–77.

3. Emerson RO, Thomas JH. Adaptive evolution in zinc finger transcription factors. PLoS Genet. 2009;5:e1000325.

4. Imbeault M, Helleboid P-Y, Trono D KRAB zinc-finger proteins contribute to the evolution of gene regulatory networks. Nature. 2017;543:550–4.

5. Wolf G, Yang P, Füchtbauer AC, Füchtbauer E-M, Silva AM, Park C, et al. The KRAB zinc finger protein ZFP809 is required to initiate epigenetic silencing of endogenous retroviruses. Genes Dev. 2015;29:538–54.

6. Ecco G, Cassano M, Kauzlaric A, Duc J, Coluccio A, Offner S, et al. Transposable Elements and Their KRAB-ZFP Controllers Regulate Gene Expression in Adult Tissues. Dev Cell. 2016;36:611–23.

7. Jacobs FMJ, Greenberg D, Nguyen N, Haeussler M, Ewing AD, Katzman S, et al. An evolutionary arms race between KRAB zinc-finger genes ZNF91/93 and SVA/L1 retrotransposons. Nature. 2014;516:242–5.

8. Najafabadi HS, Mnaimneh S, Schmitges FW, Garton M, Lam KN, Yang A, et al. C2H2 zinc finger proteins greatly expand the human regulatory lexicon. Nat Biotechnol. 2015;advance on:555–62.

9. Thomas JH, Schneider S Coevolution of retroelements and tandem zinc finger genes. Genome Res. 2011;21:1800–12.

10. Sripathy SP, Stevens J, Schultz DC. The KAP1 corepressor functions to coordinate the assembly of de novo HP1-demarcated microenvironments of heterochromatin required for KRAB zinc finger protein-mediated transcriptional repression. Mol Cell Biol. 2006;26:8623–38.

11. Macfarlan TS, Gifford WD, Agarwal S, Driscoll S, Lettieri K, Wang J, et al. Endogenous retroviruses and neighboring genes are coordinately repressed by LSD1/KDM1A. Genes Dev. 2011;25:594–607.

12. Rowe HM, Jakobsson J, Mesnard D, Rougemont J, Reynard S, Aktas T, et al. KAP1 controls endogenous retroviruses in embryonic stem cells. Nature. 2010;463:237–40.

13. Frietze S, Lan X, Jin VX, Farnham PJ. Genomic targets of the KRAB and SCAN domain-containing zinc finger protein 263. J Biol Chem. 2010;285:1393–403.

14. Yang P, Wang Y, Hoang D, Tinkham M, Patel A, Sun M, et al. A placental growth factor is silenced in mouse embryos by the zinc finger protein ZFP568. Science. 2017;356:757–9.

15. Li X, Ito M, Zhou F, Youngson N, Zuo X, Leder P, et al. A maternal-zygotic effect gene, Zfp57, maintains both maternal and paternal imprints. Dev Cell. 2008;15:547–57.

16. Bartolomei MS, Ferguson-Smith AC. Mammalian genomic imprinting. Cold Spring Harb Perspect Biol. 2011;3:1–17.

17. Quenneville S, Verde G, Corsinotti A, Kapopoulou A, Jakobsson J, Offner S, et al. In embryonic stem cells, ZFP57/KAP1 recognize a methylated hexanucleotide to affect chromatin and DNA methylation of imprinting control regions. Mol Cell. 2011;44:361–72.

18. Strogantsev R, Krueger F, Yamazawa K, Shi H, Gould P, Goldman-Roberts M, et al. Allele-specific binding of ZFP57 in the epigenetic regulation of imprinted and non-imprinted monoallelic expression. Genome Biol. 2015;16:112.

19. Takahashi N, Gray D, Strogantsev R, Noon A, Delahaye C, Skarnes WC, et al. ZFP57 and the Targeted Maintenance of Postfertilization Genomic Imprints. Cold Spring Harb Symp Quant Biol. 2015;80:177–87.

20. Takahashi N, Coluccio A, Thorball CW, Planet E, Shi H, Offner S, et al. ZNF445 is a primary regulator of genomic imprinting. Genes Dev. 2019;33:49–54.

21. Inoue A, Jiang L, Lu F, Suzuki T, Zhang Y Maternal H3K27me3 controls DNA methylation-independent imprinting. Nature. 2017;547:419–24.

22. Boonen SE, Mackay DJG, Hahnemann JMD, Docherty L, Grønskov K, Lehmann A, et al. Transient neonatal diabetes, ZFP57, and hypomethylation of multiple imprinted loci: a detailed follow-up. Diabetes Care. 2013;36:505–12.

23. Mackay DJG, Callaway JL a, Marks SM, White HE, Acerini CL, Boonen SE, et al. Hypomethylation of multiple imprinted loci in individuals with transient neonatal diabetes is associated with mutations in ZFP57. Nat Genet. 2008;40:949–51.

24. Riso V, Cammisa M, Kukreja H, Anvar Z, Verde G, Sparago A, et al. ZFP57 maintains the parent-of-origin-specific expression of the imprinted genes and differentially affects non-imprinted targets in mouse embryonic stem cells. Nucleic Acids Res. 2016;44:8165–78.

25. Berrens R V., Andrews S, Spensberger D, Santos F, Dean W, Gould P, et al. An endosiRNA-Based Repression Mechanism Counteracts Transposon Activation during Global DNA Demethylation in Embryonic Stem Cells. Cell Stem Cell. 2017;21:694–703.e7.

26. Karimi MM, Goyal P, Maksakova IA, Bilenky M, Leung D, Tang JX, et al. DNA methylation and SETDB1/H3K9me3 regulate predominantly distinct sets of genes, retroelements, and chimeric transcripts in mESCs. Cell Stem Cell. 2011;8:676–87.

27. Nichols J, Jones K Derivation of Mouse Embryonic Stem (ES) Cell Lines Using Small-Molecule Inhibitors of Erk and Gsk3 Signaling (2i). Cold Spring Harb Protoc. 2017;2017:pdb.prot094086.

28. Proudhon C, Duffié R, Ajjan S, Cowley M, Iranzo J, Carbajosa G, et al. Protection against De Novo Methylation Is Instrumental in Maintaining Parent-of-Origin Methylation Inherited from the Gametes. Mol Cell. 2012;47:909–20.

29. Hendrickson PG, Doráis J a, Grow EJ, Whiddon JL, Lim J-W, Wike CL, et al. Conserved roles of mouse DUX and human DUX4 in activating cleavage-stage genes and MERVL/HERVL retrotransposons. Nat Genet. 2017;49:925–34.

30. Iaco A De, Planet E, Coluccio A, Verp S, Duc J, Trono D Europe PMC Funders Group Europe PMC Funders Author Manuscripts A family of double-homeodomain transcription factors regulates zygotic genome activation in placental mammals. 2017;49:941–5.

31. Wang L, Zhang J, Duan J, Gao X, Zhu W, Lu X, et al. Programming and inheritance of parental DNA methylomes in mammals. Cell. 2014;157:979–91.

32. Smith ZD, Shi J, Gu H, Donaghey J, Clement K, Cacchiarelli D, et al. Epigenetic restriction of extraembryonic lineages mirrors the somatic transition to cancer. Nature. 2017;549:543–7.

33. Kobayashi H, Sakurai T, Imai M, Takahashi N, Fukuda A, Yayoi O, et al. Contribution of intragenic DNA methylation in mouse gametic DNA methylomes to establish oocyte-specific heritable marks. PLoS Genet. 2012;8:e1002440.

34. Smallwood SA, Tomizawa S-I, Krueger F, Ruf N, Carli N, Segonds-Pichon A, et al. Dynamic CpG island methylation landscape in oocytes and preimplantation embryos. Nat Genet. 2011;43:811–4.

35. Stewart KR, Veselovska L, Kelsey G Establishment and functions of DNA methylation in the germline. Epigenomics. 2016;8:1399–413.

36. Blahnik KR, Dou L, Echipare L, Iyengar S, O’Geen H, Sanchez E, et al. Characterization of the contradictory chromatin signatures at the 3’ exons of zinc finger genes. PLoS One. 2011;6:e17121.

37. Valle-García D, Qadeer ZA, McHugh DS, Ghiraldini FG, Chowdhury AH, Hasson D, et al. ATRX binds to atypical chromatin domains at the 3' exons of zinc finger genes to preserve H3K9me3 enrichment. Epigenetics. 2016;11:398–414.

38. Dodge JE, Kang Y, Beppu H, Lei H Histone H3-K9 Methyltransferase ESET Is Essential for Early Development Histone H3-K9 Methyltransferase ESET Is Essential for Early Development. Mol Cell Biol. 2004;24:2478–86.

39. Messerschmidt DM, de Vries W, Ito M, Solter D, Ferguson-Smith A, Knowles BB. Trim28 is required for epigenetic stability during mouse oocyte to embryo transition. Science. 2012;335:1499–502.

40. Lehnertz B, Ueda Y, Derijck AAHA, Braunschweig U, Perez-Burgos L, Kubicek S, et al. Suv39h-mediated histone H3 lysine 9 methylation directs DNA methylation to major satellite repeats at pericentric heterochromatin. Curr Biol. 2003;13:1192–200.

41. Liu Y, Olanrewaju YO, Zhang X, Cheng X DNA recognition of 5-carboxylcytosine by a Zfp57 mutant at an atomic resolution of 0.97 Å. Biochemistry. 2013;52:9310–7.

42. Friedli M, Trono D The developmental control of transposable elements and the evolution of higher species. Annu Rev Cell Dev Biol. 2015;31:429–51.

43. Chuong EB, Rumi M a K, Soares MJ, Baker JC. Endogenous retroviruses function as species-specific enhancer elements in the placenta. Nat Genet. 2013;45:325–9.

44. Rowe HM, Kapopoulou A, Corsinotti A, Fasching L, Macfarlan TS, Tarabay Y, et al. TRIM28 repression of retrotransposon-based enhancers is necessary to preserve transcriptional dynamics in embryonic stem cells. Genome Res. 2013;23:452–61.

45. Thompson PJ, Macfarlan TS, Lorincz MC. Long Terminal Repeats: From Parasitic Elements to Building Blocks of the Transcriptional Regulatory Repertoire. Mol Cell. 2016;62:766–76.

