## Supplementary material for "Epigenetic regulation of unique genes and repetitive elements by the KRAB zinc finger protein ZFP57": Fig. S1-S11

**Supplementary Materials**

Materials and Methods

**ES cell culture**

Embryonic Stem cells were derived in ground state pluripotency conditions as described previously [1]. Briefly, morula stage embryos were collected at E2.5 and cultured for 2 days in KSOM with inhibitors of Erk and Gsk3 (2i) signaling pathways (PD0325901 and CHIR99021). The trophectoderm was then removed using complement immunosurgery and used for genotyping. The inner cell mass was cultured in N2B27 media supplemented with 2i and LIF until establishment of stable ES cell cultures. Each ES cell line was characterized by staining for alkaline phosphatase activity and expression of Oct-4 (SantaCruz, sc8628) and SSEA1 (Millipore, MAB4301) markers. Finally, each newly established line was karyotyped and screened for mycoplasma contamination prior to use in downstream experiments. ES cells were cultured under 2i/LIF conditions up to a maximum of 15 passages, matching the passage number for WT and KO cells for each experiment.

**DNA and RNA extraction**

DNA was extracted using tale lysis buffer (50mM Tris-Cl pH8.5; 5mM EDTA; 200mM Nacl; 0.25% SDS) and 1 µg/ml Proteinase K for 1 hr. The mixture was purified with phenol/chloroform extraction and precipitated with isopropanol. RNA was extracted using Trizol (SIGMA) according to manufacturer’s protocol. RNA/DNA was quantified using Picodrop and quality assayed by gel electrophoresis.

For RNA-seq library preparation, total Trizol extracted RNA (1 µg) was ribodepleted using NEBNext rRNA depletion kit (NEB, E6310L) according to manufacturer’s protocol and purified using Agencourt RNAClean XP beads (Beckman Coulter, A63987). RNA-seq library was prepared using ScripSeq v2 kit (Illumina, SSV21124) with 10 cycles of amplification. Successful ribodepletion was confirmed by post-sequencing QC library analysis. All sequencing was performed with pair-end reads using Illumina HiSeq platforms.

For analysis, RNA-seq reads were first quality checked with FastQC (version 0.11.5), followed by adapter and base quality trimming with TrimGalore (version 0.2.7). Reads post-QC were mapped to mouse reference genome (mm10) with STAR (version 2.5.2, --alignEndsType EndToEnd --outFilterMultimapNmax 300 --outSAMmultNmax 1 --outMultimapperOrder Random) [2]. Expression of unique genes and transposable elements were quantified with featureCounts [3], which were subsequently converted into FPKM (fragments per kilobase per million mapped reads), representing gene expression levels. Differential expression analysis was performed with DESeq2 [4]. Annotations for RefSeq genes and transposable elements were downloaded from the UCSC genome browser [5]. Visualization of RNA-seq data was performed with RSeQC (version 2.4) [6] to convert data into BigWig format and WashU Epigenome Browser [7] were utilized to visualize the data.

**ChIP-seq**

ChIP sequencing libraries from wild type and ZFP57 KO ES cells were prepared as described previously [8]. The following antibodies were used for immunoprecipitation: ZFP57 (abcam ab45341, lot 754226); KAP1 (abcam ab10483, lot J0809) and H3K9me3 (abcam, ab8898, lot GR285794-1). The libraries were prepared using NEBNext UltraII kit (E7645).

Similar QC steps including FastQC and TrimGalore were performed on ChIP-seq reads. Reads post-QC were aligned to mouse reference genome (mm10) with BWA (version 0.6.2) with pair-end reads mode [9]. Potential PCR duplicates were removed with Picard tools ("MarkDuplicates" function). Peak calling was performed with MACS2 (version 2.1.0) [10] with broad peak option using only uniquely aligned reads (MAPQ>10). ZFP57 peaks were normalized to ChIP-seq library in ZFP57 mutant ES cells, and KAP1, H3K9me3 peaks were normalized to their corresponding input control. Peaks called in at least two of the three WT biological replicates were treated as high-confidence peaks, and were used for subsequent analyses. Enrichment analysis on genomic features was performed with bedtools2 software (version 2.27.0) to generate intersection, identify closest unique genes/repetitive elements and to generate control shuffled regions. Gene promoters were defined as 1kb ± transcription start sites (TSSs).

Repeat content of ZFP57 peaks was determined based on genomic overlaps with repeats annotated in UCSC Genome Browsers (http://genome.ucsc.edu/) [5]. Peaks were characterized into repetitive and unique according to two factors, 1) if the fraction of overlap with repeat is >10%, and 2) location of ZFP57 motif – whether it reside in the repeat or the unique proportion of the peak. Heat maps of ChIP-seq results were plotted using R Bioconductor package "Genomation" [11]. Visualization tracks were generated using bedtools2 genomecov (version 2.27.0, -pc -bga -scale) with scaling factor being per million ($10^6$) the number aligned reads. Visualization of ChIP-seq data was performed with WashU Epigenome Browser [7] and IGB viewer [12].

**Bisulfite treatment and pyrosequencing**

Purified DNA was bisulfite converted using SIGMA Imprint (MOD50) DNA modification kit and subjected to PCR and pyrosequencing analysis as described previously [8].

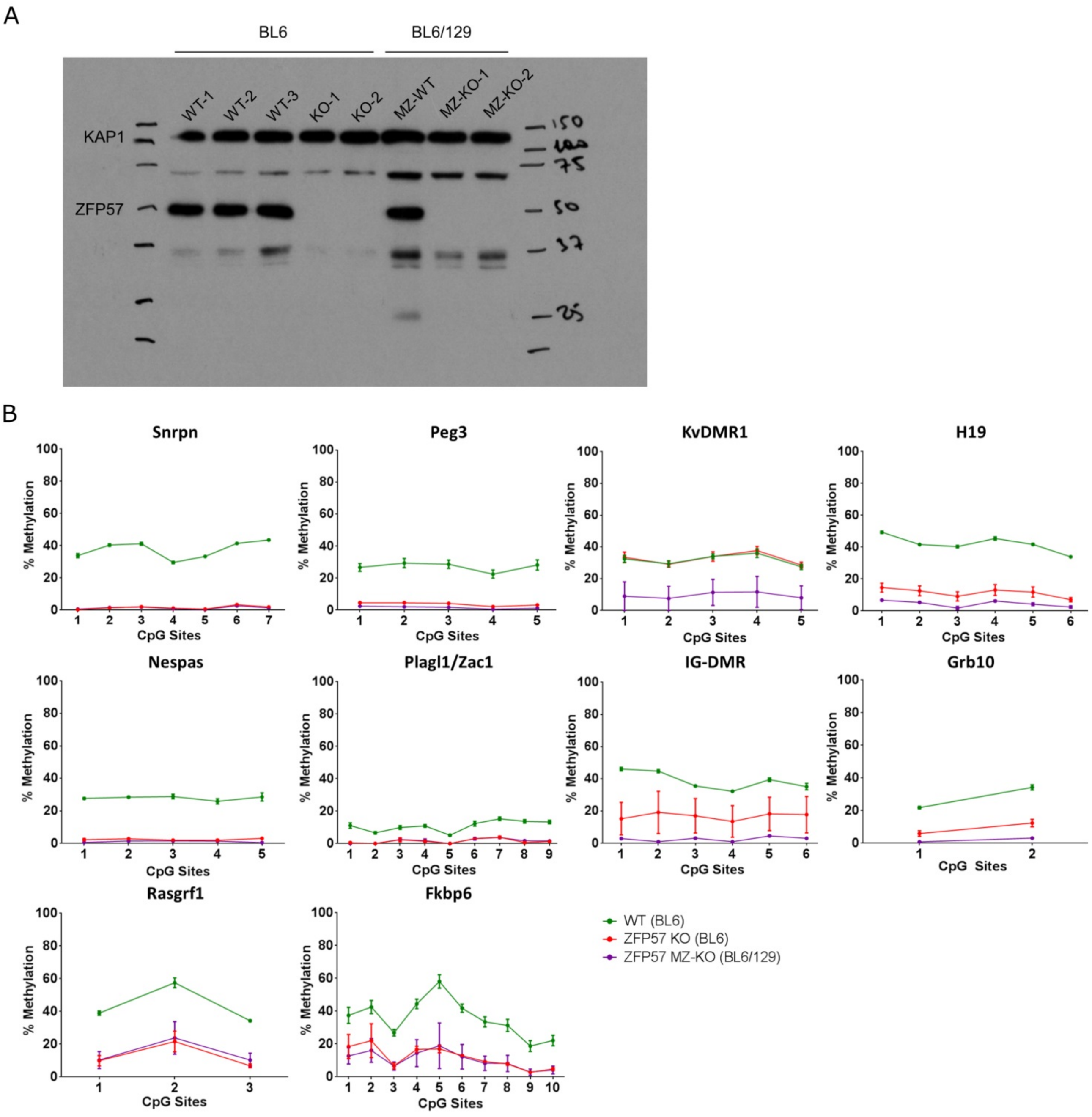


**Fig. S1. Analysis of blastocyst derived ZFP57 null ES cells.** (A) Western Blot analysis confirming absence of detectable ZFP57 protein. (B) Bisulfite pyrosequencing of ICRs in WT, ZFP57 KO and ZFP57 MZ-KO ES cells.

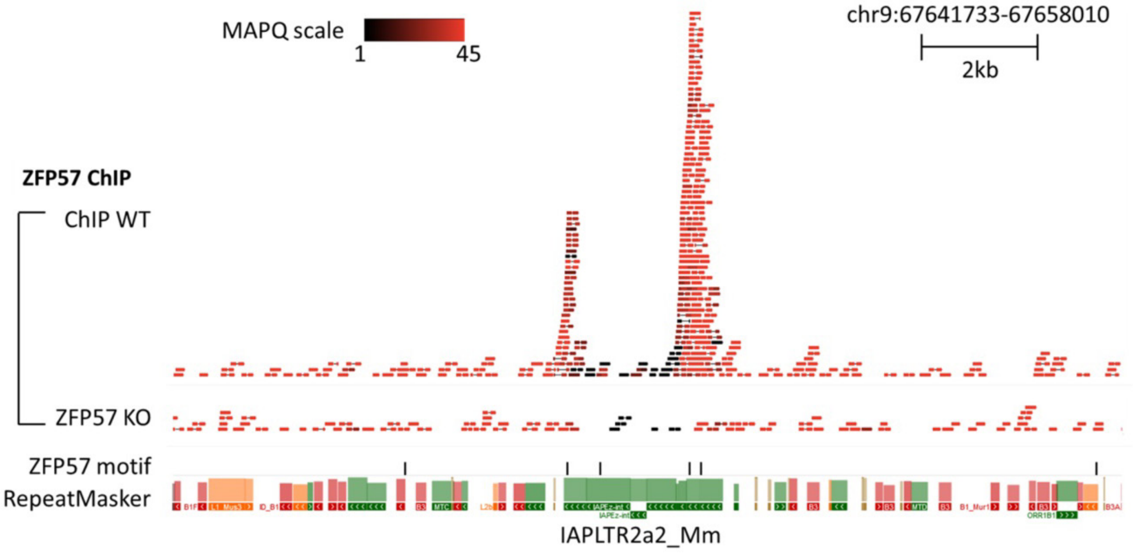


**Fig. S2.** IGB viewer snapshot showing representative ZFP57 peak at IAPLTR2a2_Mm retroelements, which harbours the most number of ZFP57 peaks. Pair-end sequencing, together with sequence divergence of the retroelements allowing us to map ChIP-seq reads to individual transposable elements in the genome. The colour scale indicates the mapping quality score (MAPQ) for each read pair. MAPQ = $-10 log_{10} P$ where $P$ is the probability that true alignment belongs elsewhere.

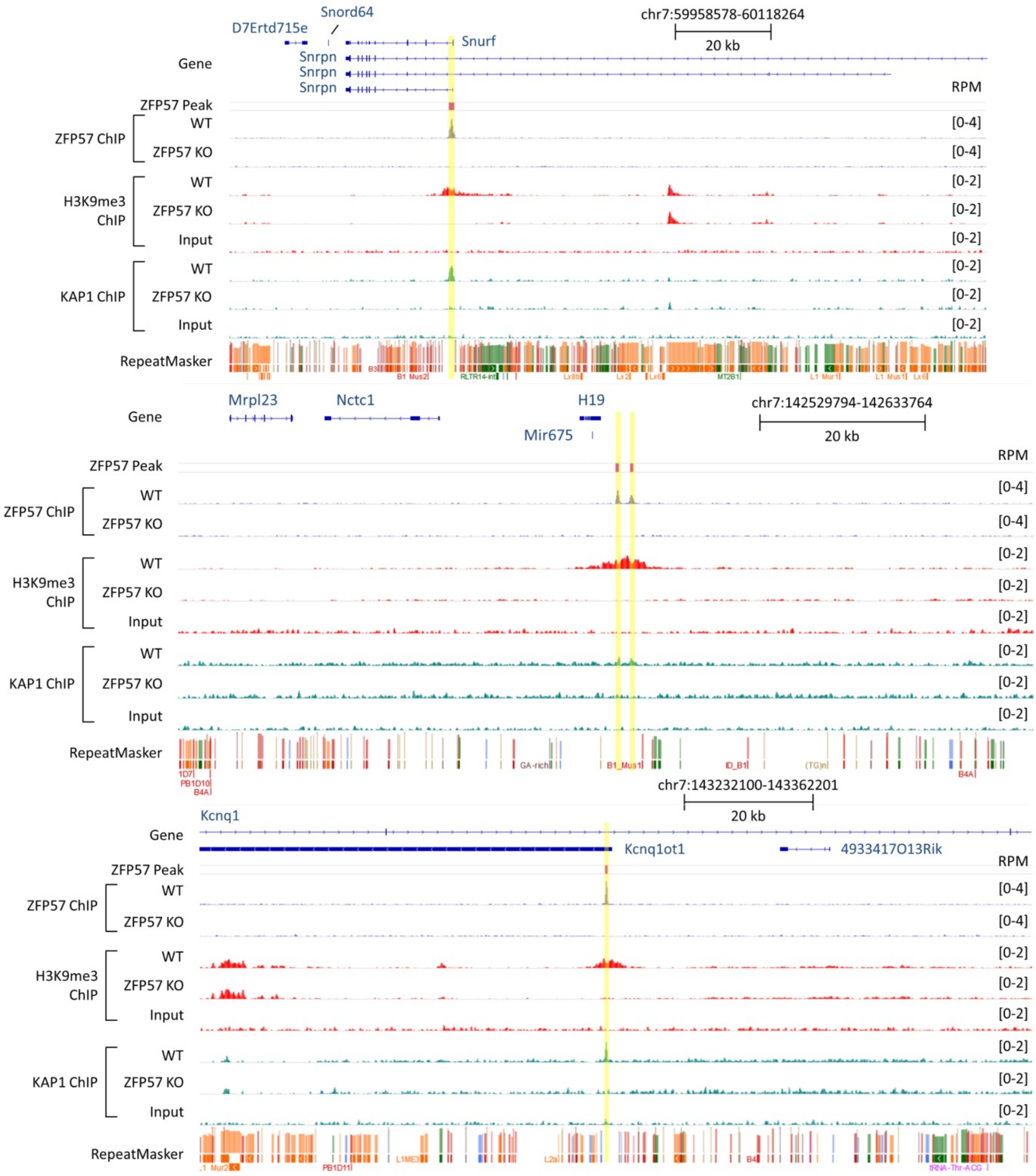


**Fig. S3.** Genome browser screenshots for Snrpn, H19 and Kcnq1ot1 imprinted loci.

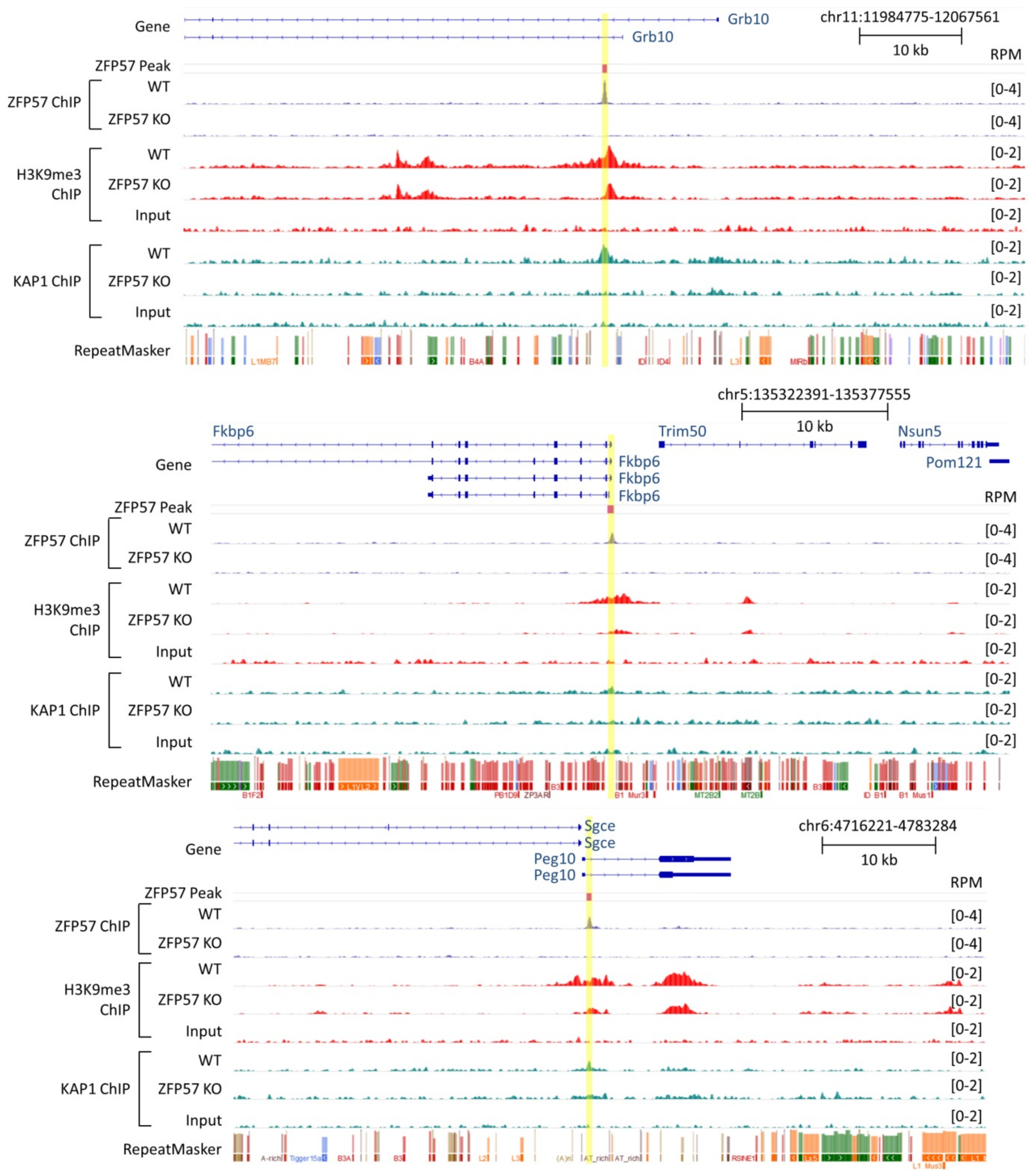


**Fig. S4.** Genome browser screenshots for Grb10, Fkbp6 and Peg10 imprinted loci.

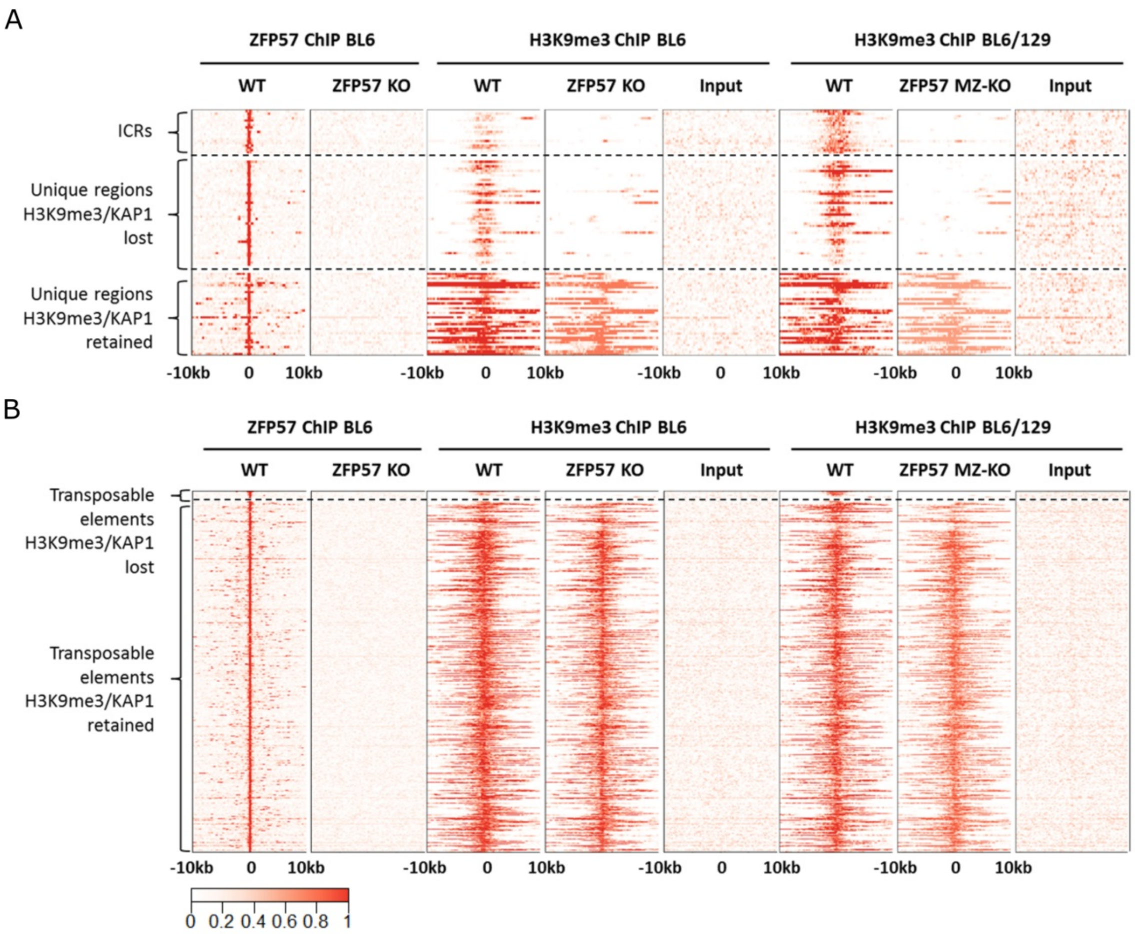


**Fig. S5.** Heat maps showing signal enrichment of H3K9me3 at ZFP57 binding sites in ZFP57 KO and MZ-KO ES cells.

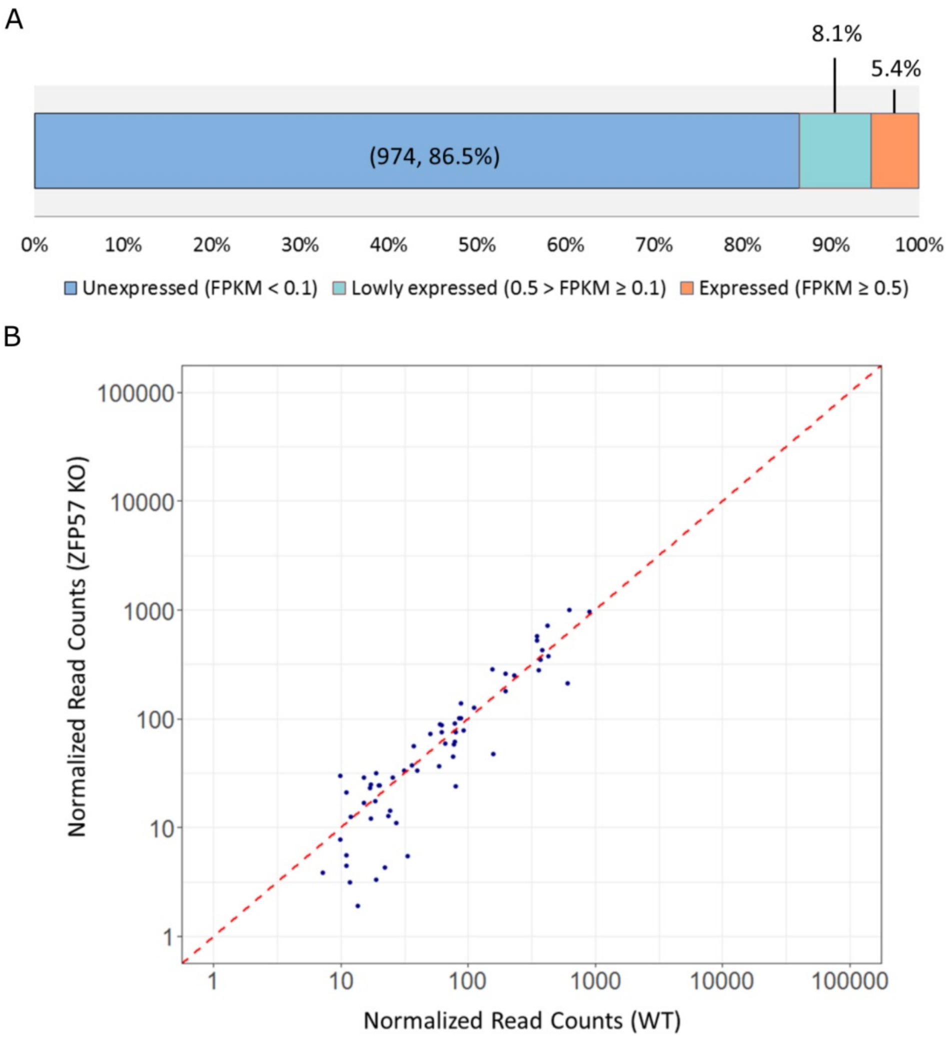


**Fig. S6. Transposable elements bound by ZFP57 are not reactivated in ZFP57 KO ES cells.** (A) Expression status of ZFP57 bound transposable elements in WT and ZFP57 mutants. (B) Scatterplot showing no transcriptional changes of TEs (FPKM ≥ 0.1 in WT) in ZFP57 KO cells versus WT.

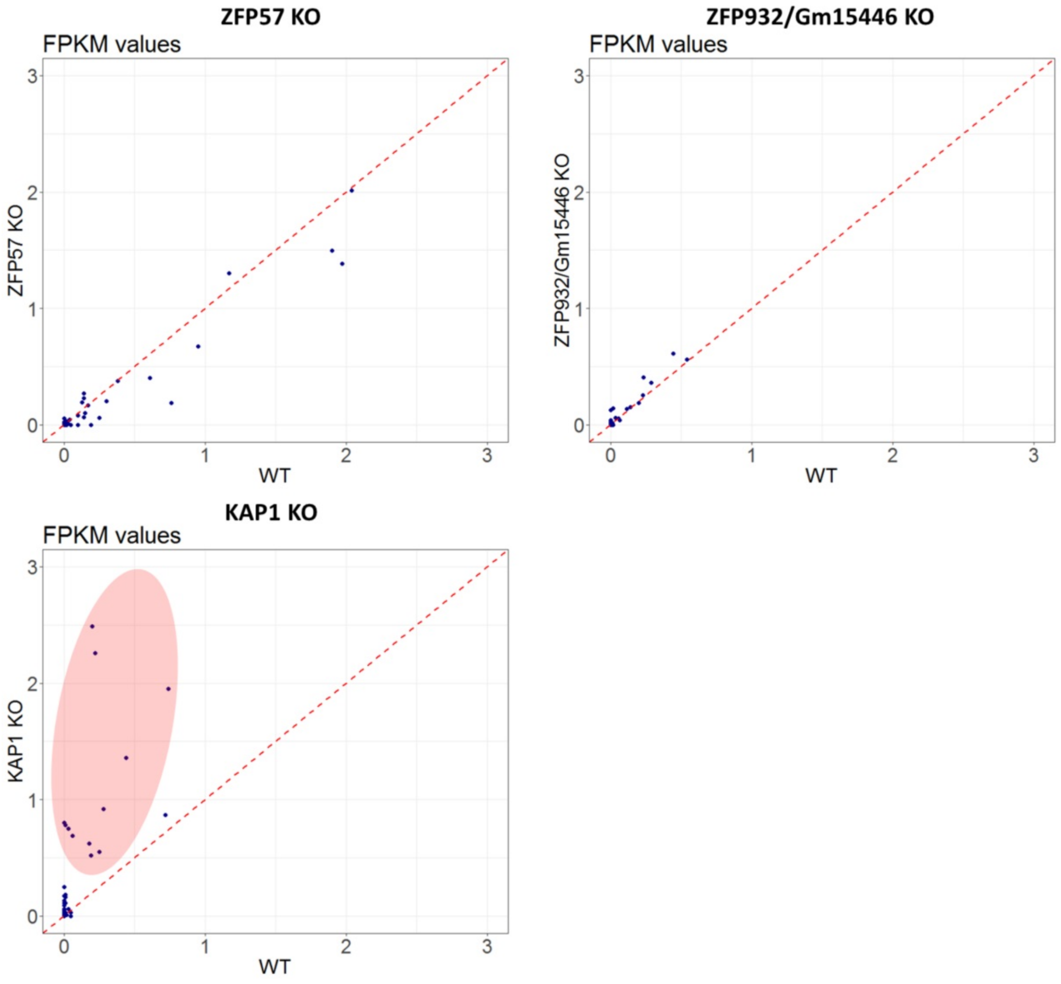


**Fig. S7. Analysis of expression of transposable elements jointly bound by ZFP57, ZFP932 and/or Gm15446 in different mutant ES cells.** Top left – ZFP57 KO, Top right – ZFP932/Gm15446 DKO, Bottom left – KAP1 KO. The IAP retroelement in vicinity of Bglap3 locus is not shown in the ZFP932/Gm15446 DKO as its expression far exceeds other jointly bound TEs [13].

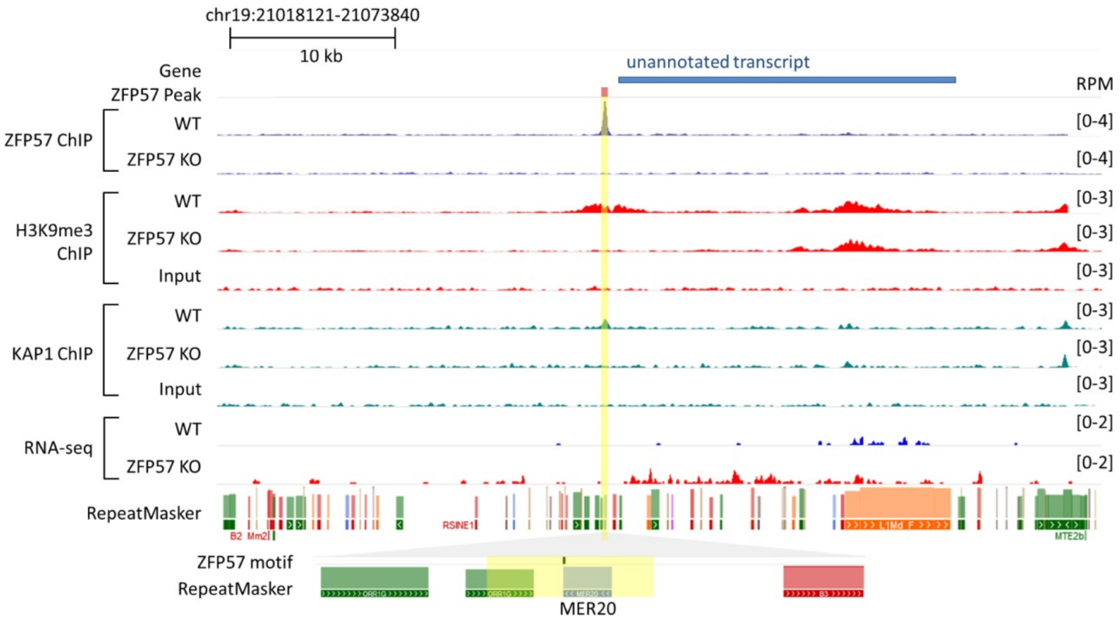


**Fig. S8.** Genome browser screenshot showing loss of ZFP57 binding at MER20 DNA transposon resulting in loss of KAP1 and H3K9me3 and upregulation of an unannotated transcript.

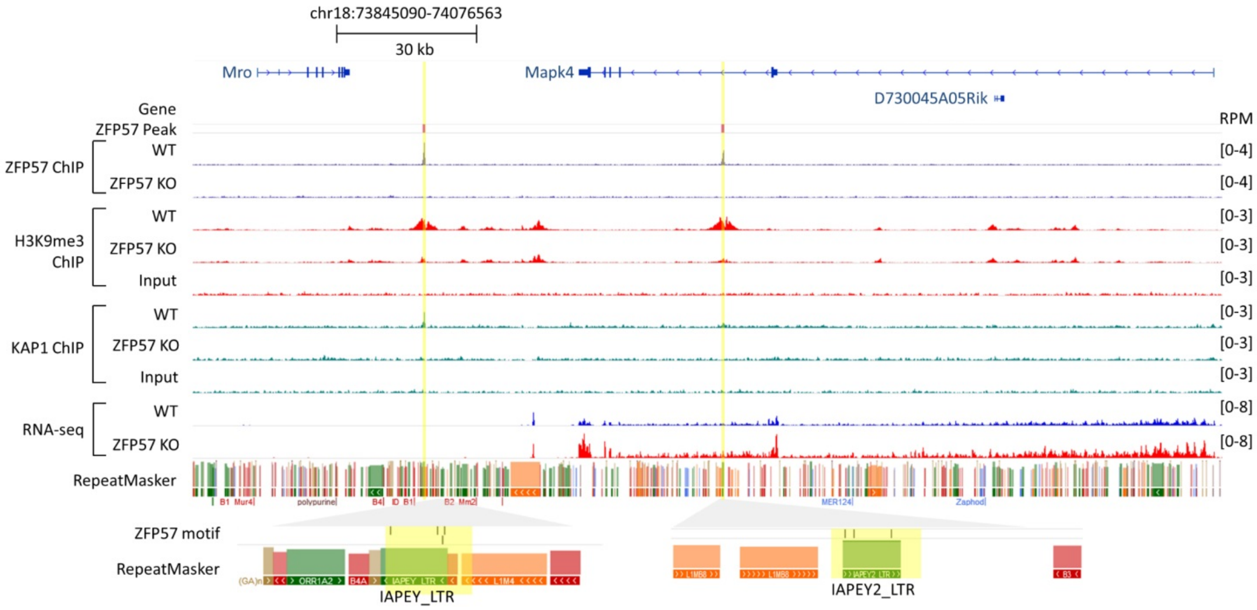


**Fig. S9.** Genome browser screenshot showing loss of KAP1 and H3K9me3 in solo LTRs located within and adjacent to Mapk4 gene, and an increase in Mapk4 levels in ZFP57 KO ES cells.

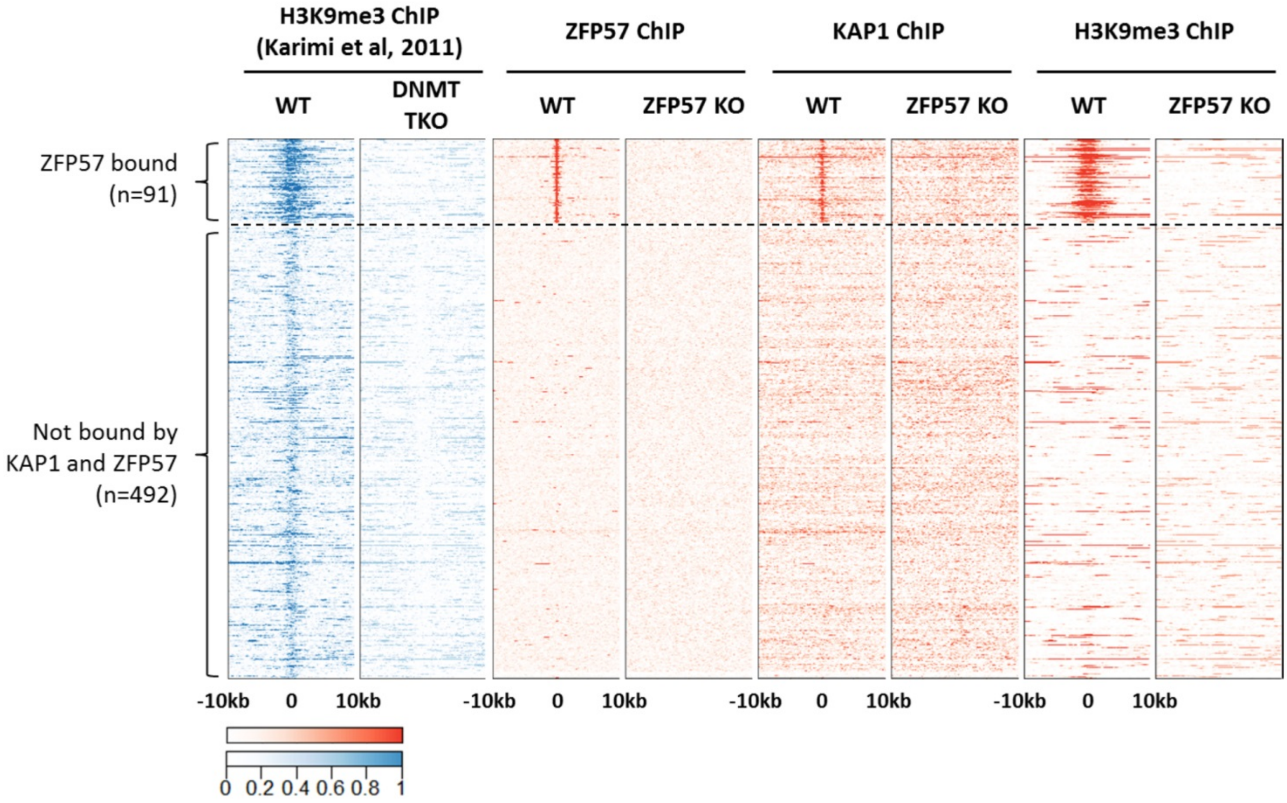


**Fig. S10. Heat maps of DNA methylation dependent H3K9me3 enriched regions in WT and DNMT TKO cells** [14]**.** Enrichment of ZFP57, KAP1 and H3K9me3 in ZFP57 KO ES cells at the same regions.

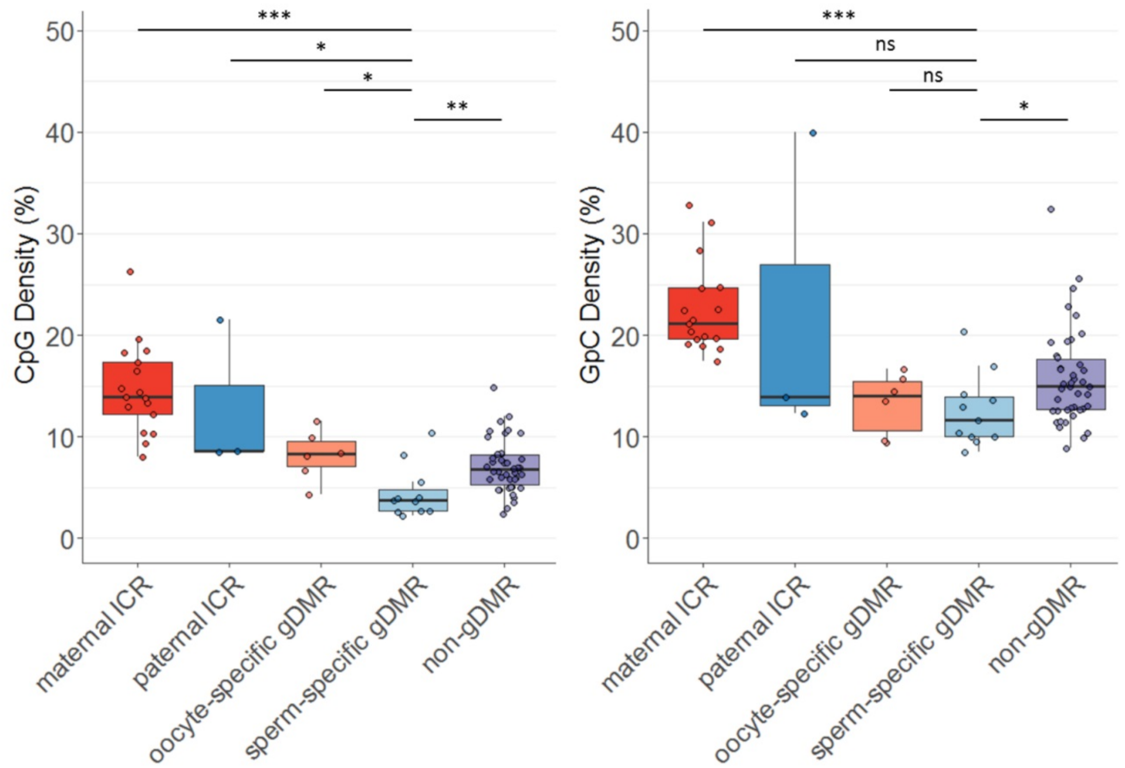


**Fig. S11.** Box and whisker plot, showing CpG density (left) and GpC density (right) for five different categories of ZFP57 targets – maternal ICRs, paternal ICRs, oocyte-specific gDMR, sperm-specific gDMR and non-gDMRs methylated in both germlines. ns – not significant, * *P* < 0.05, ** *P* < 0.01, *** *P* < 0.001; Mann-Whitney *U* Test.

### References


1. Nichols J, Jones K. Derivation of Mouse Embryonic Stem (ES) Cell Lines Using Small-Molecule Inhibitors of Erk and Gsk3 Signaling (2i). Cold Spring Harb Protoc. 2017;2017:pdb.prot094086.
2. Dobin A, Davis C a, Schlesinger F, Drenkow J, Zaleski C, Jha S, et al. STAR: ultrafast universal RNA-seq aligner. Bioinformatics. 2013;29:15–21.
3. Liao Y, Smyth GK, Shi W. FeatureCounts: An efficient general purpose program for assigning sequence reads to genomic features. Bioinformatics. 2014;30:923–30.
4. Love MI, Huber W, Anders S. Moderated estimation of fold change and dispersion for RNA-seq data with DESeq2. Genome Biol. 2014;15:550.
5. Kent WJ, Sugnet CW, Furey TS, Roskin KM, Pringle TH, Zahler AM, et al. The Human Genome Browser at UCSC. Genome Res. 2002;12:996–1006.
6. Wang L, Wang S, Li W. RSeQC: quality control of RNA-seq experiments. Bioinformatics. 2012;28:2184–5.
7. Zhou X, Li D, Zhang B, Lowdon RF, Rockweiler NB, Sears RL, et al. Epigenomic annotation of genetic variants using the Roadmap Epigenome Browser. Nat Biotechnol. 2015;33:345–6.
8. Strogantsev R, Krueger F, Yamazawa K, Shi H, Gould P, Goldman-Roberts M, et al. Allele-specific binding of ZFP57 in the epigenetic regulation of imprinted and non-imprinted monoallelic expression. Genome Biol. 2015;16:112.
9. Li H, Durbin R. Fast and accurate short read alignment with Burrows-Wheeler transform. Bioinformatics. 2009;25:1754–60.
10. Feng J, Liu T, Qin B, Zhang Y, Liu XS. Identifying ChIP-seq enrichment using MACS. Nat Protoc. 2012;7:1728–40.
11. Akalin A. Genomation a Toolkit for Annotation and Visualization of Genomic Data. 2013;1–15.
12. Freese NH, Norris DC, Loraine AE. Integrated genome browser: Visual analytics platform for genomics. Bioinformatics. 2016;32:2089–95.
13. Ecco G, Cassano M, Kauzlaric A, Duc J, Coluccio A, Offner S, et al. Transposable Elements and Their KRAB-ZFP Controllers Regulate Gene Expression in Adult Tissues. Dev Cell. 2016;36:611–23.
14. Karimi MM, Goyal P, Maksakova IA, Bilenky M, Leung D, Tang JX, et al. DNA methylation and SETDB1/H3K9me3 regulate predominantly distinct sets of genes, retroelements, and chimeric transcripts in mESCs. Cell Stem Cell. 2011;8:676–87.
